# Sequential assembly *in vivo* of the bacteriophage P22 genome translocation channel

**DOI:** 10.64898/2026.09.03.749273

**Authors:** Huaxin Yu, Jian Yue, Taehyun Park, Chunyan Wang, Muyuan Chen, Jun Liu, Ian J. Molineux

## Abstract

Bacteriophage P22 ejects four internal proteins into infected cells to assemble a channel for genome translocation across the cell envelope. However, how these proteins cooperate to build the trans-envelope channel remains enigmatic. Using *in situ* single-particle cryogenic electron microscopy, we determined near-atomic *in vivo* structures of P22 infection intermediates, revealing how the tail hub protein gp10 templates the sequential assembly of three internal proteins gp7*, gp20 and gp16 into a ∼50-nm trans-envelope channel that extends from the virion into the host cytoplasm. Functional analyses explain why phage DNA can be detected in supernatant fluids during infection by channel-defective mutants. Our structures also provide a mechanistic basis for the long-standing observation that a P22 virion lacking gp16 can be complemented, extra-cytoplasmically, by a co-infecting virion lacking gp20, thereby allowing the genome in the gp16-deficient particle to enter the host cytoplasm.

## Introduction

Bacteriophages have evolved sophisticated mechanisms to adsorb and infect bacteria with high specificity. They often employ structurally distinct tail machines, categorized into three major families: myophages possess a rigid contractile tail, siphophages a flexible non-contractile tail, and podophages a short stubby tail. These distinct tail machines are each capable of breaching the multiple barriers presented by both Gram-positive and Gram-negative bacterial envelopes. Multiple approaches have been utilized to investigate the tail machines during phage infection. In particular, cryo-electron tomography (cryo-ET) has emerged as a powerful tool for studying conformational changes during infection of *Prochlorococcus marinus* phage P-SSP7(*1, 2*), *Caulobacter crescentus* phage ΦCb13(*3*), *E. coli* phages P1(*4*), λ(*5*), T7(*6*) and T4(*7*), *S. aureus* Φ812(*8*), *Bacillus subtilis* phage Φ29(*9*), *Salmonella* Typhimurium phages SP6(*10*) and P22(*11*), and *Pseudomonas aeruginosa* phage JBD30(*12*). Recent cryo-electron microscopy (cryo-EM) studies of *E. coli* phages T7(*13, 14*), SU10(*15*), T5(*16*) and DT57C(*17*), λ(*18*) and *Pseudomonas* phage E217(*19*) reveal striking conformational changes triggered by receptor binding *in vitro*. However, despite these advances, how phages breach the bacterial envelope to establish genome translocation during productive infections *in vivo* remain unresolved.

P22, a stubby-tailed podophage that infects smooth strains of *S.* Typhimurium has long served as a model system for studying viral assembly and infection mechanisms(*20, 21*). Recent cryo-EM structures provided near-atomic details of mature P22(*22, 23*), including its capsid (gp5), portal (gp1), tail adaptor (gp4), tail hub (gp10), tailspike (gp9) and needle (gp26). Gp26 and three internal virion proteins (gp7, gp16, and gp20) are called E proteins because they are selectively lost from phage particles that could be desorbed from infected cells and the proteins were therefore assumed to be ejected into the host cell(*24*). It was later shown that they are indeed located inside the infected cell(*11, 23*). The E proteins are all essential for phage infection(*20*). Gp7*, the proteolytically processed, active form of gp7(*25, 26*) appears to form the extracellular channel bridging the tail hub and the host cell surface; gp20 and gp16 then extend the channel across the cell envelope(*11*). It is not understood how the E proteins cooperate in assembling the channel. It was also reported 50 years ago that a mutant virion lacking gp16, which is unable to eject its DNA into the cell cytoplasm, can be complemented by co-infection with a mutant virion lacking gp20(*27, 28*). The latter mutant particle is also unable to eject its DNA into the cell cytoplasm. Further, this extra-cytoplasmic complementation is unidirectional – only the genome from the gp16^-^ particle initiates phage development. No molecular understanding of these observations has been reported.

Here, we develop an *in situ* single-particle cryo-EM approach to visualize *Salmonella* minicells infected by wild-type P22 and mutant particles at far higher resolution than has hitherto been possible. This has enabled determining, at near-atomic resolution, intermediate and the final structure of an *in vivo* phage-induced channel that spans the outer envelope. Our study also provides the first structural explanation for the remarkable extra-cytoplasmic complementation that occurs through proteins delivered from co-infecting defective virions.

## Results

### Visualizing P22 infection using *in situ* cryo-EM and cryo-ET

Among the well-characterized gene products of P22 (**Fig. 1a**), three E proteins – gp7*, gp20, and gp16 are stored inside mature virions and are ejected into the infected host cell where they assemble into a trans-envelope channel between the gp10 tail hub and host cytoplasm. Part of the channel structure was previously obtained at ∼3 nm resolution using cryo-ET(*11*), insufficient for a mechanistic understanding of how its assembly occurs.

**Figure 1.**
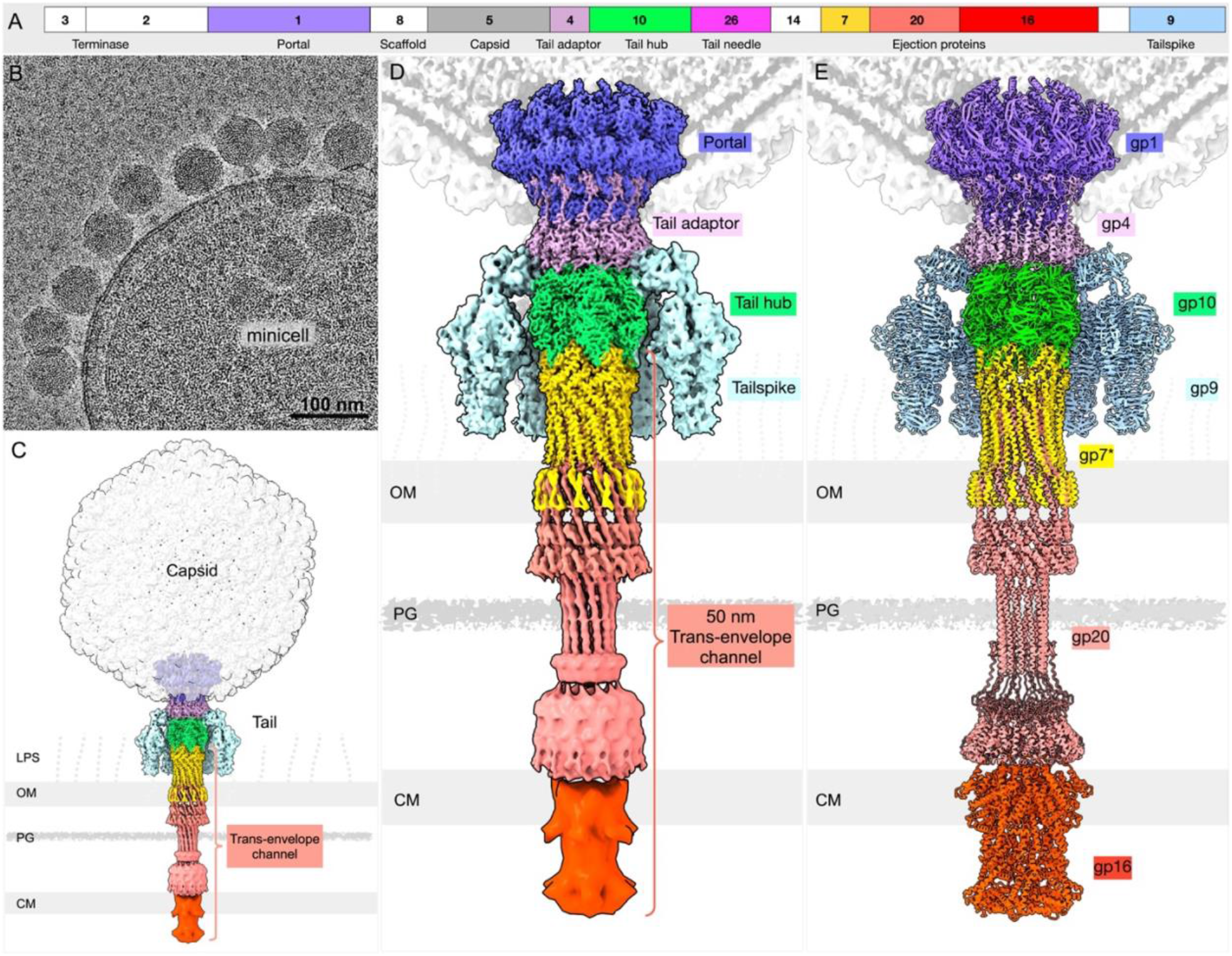
*In situ* structure of a P22 infection intermediate. (**a**) Organization of P22 genes encoding structural proteins. Non-virion genes are shown as white boxes; several genes lying between genes *16* and *9* are omitted for clarity. The same color scheme is used throughout subsequent figures. (**b**) Representative cryo-EM micrograph of a P22-infected minicell. (**c**) Overall *in situ* structure of a P22 infection intermediate, revealing a DNA ejection channel extending across the cytoplasmic membrane. (**d**) Enlarged view of the extended tail complex during infection, including the portal, tail adaptor, hub, tailspike, and three ejection proteins that form a 50-nm trans-envelope channel. (**e**) Atomic model of the extended tail complex during infection, including gp1, gp4, gp10, with gp7*, gp20 and gp16 forming the channel based on the integration of cryo-EM structures and density-constrained AlphaFold3 predictions. Six copies of gp16 were used in the AlphaFold3 prediction, as this number best fits the cryo-EM density.

To improve resolution, we employed an advanced cryo-ET data processing pipeline implemented in EMAN2 that integrates tilt series alignment, tomogram reconstruction and subtomogram averaging into a streamlined workflow(*29*). This automated pipeline enabled us to efficiently analyze 731 tilt series of *Salmonella* minicells infected by wild-type P22 **(fig. S1)**. More than 40,000 adsorbed particles were automatically picked for structure determination. Following determination of phage tail orientation and then multiple rounds of classification and focused refinement, we resolved the extracellular channel structure to an overall resolution 8.7 Å, enabling visualization of the α-helical backbone and revealed the attachment of gp7* to the tail hub (**fig. S1c**). The outermost portion of the periplasmic channel was resolved to 16 Å resolution (**fig. S1d**), whereas the region adjacent to the cytoplasmic membrane was less well resolved.

To further improve the structural details of the genome translocation channel, we developed an innovative *in situ* single-particle cryo-EM approach to analyze P22-infected *Salmonella* minicells. Single-particle cryo-EM has traditionally been used to determine structures of purified phage particles under cell-free conditions but, to the best of our knowledge, it has not previously been applied successfully to visualize phage infection intermediates in their native cellular environment at high resolution. To achieve sufficient cryo-EM image quality for *in situ* structure determination, we optimized sample preparation by enriching for tiny minicells through a two-step differential centrifugation protocol(*6, 11*), minimizing vitreous ice thickness, and optimizing both the phage-to-cell ratio and sample concentration. Importantly, high-throughput cryo-EM data acquisition was performed using SerialEM(*30*) with a customized multishot script(*31*) designed to specifically target phage-infected minicells (**Fig. 1b)**. From 13,298 micrographs, a total of 322,345 capsid particles were picked automatically using Topaz(*32*) and subjected to 3D classification in RELION(*33*) to localize the phage tail **(fig. S2)**. Tailed particles were exported to cryoSPARC(*34*) to perform iterative rounds of focused refinement, yielding a structure of the P22 tail machine bound to a cell at high resolution (**Fig. 1c, d; fig. S2-6**). The extended tail structure spanning from the gp10 hub through the extracellular space and outer membrane to the peptidoglycan layer was resolved between 3.1 and 5.2 Å (**fig. S5)**, several-fold higher than that achieved using cryo-ET and subtomogram averaging (**fig. S1**). Notably, focused classification revealed a stump-like density spanning the cytoplasmic membrane (**Fig. 1c, d).** This density is shown below corresponds to gp16 (**fig. S4b)**.

By integrating cryo-EM density with AlphaFold3 predictions, we built a near-complete atomic model of the P22 infection intermediate (**Fig. 1e, Movie S1**). As expected(*24*), after E protein ejection the entire barrel domain of the gp1 portal has collapsed, whereas the adaptor protein gp4 and the gp9 tailspike are structurally indistinguishable from their counterparts in mature, cell-free P22(*22, 23*) (**fig. S6b, c; fig. S7a, b**). In contrast, the gp10 hub undergoes pronounced conformational arrangements, and gp7* (residues 29–229; residues 1–20 are proteolytically removed before virion assembly) and gp20 (residues 1–247) are resolved for the first time as channel-forming proteins (**fig. S6d–l**). In mature virions both proteins are packaged in a disordered arrangement, at least in part in the portal barrel and lumen, extending through the tail lumen to the gp10 hub(*22, 23*). These observations show that, during infection, release of the internal E proteins from the capsid triggers collapse of the portal barrel while simultaneously promoting tail extension and assembly of the trans-envelope channel.

### Gp7* interacts with the gp10 tail hub in two distinct conformations

During infection, gp10 serves as the foundation for assembly of the gp7* extracellular channel(*23*). *In situ* single-particle cryo-EM reveals atomic level details of the gp10–gp7* complex in which residues 90–126 of two copies of the gp7* dodecamer form a fingertip that fills a wedge between two adjacent gp10 subunits (**Fig. 2a**). The adjacent gp7* pair adopts two distinct conformations: one (gp7*-o) is located outside while the other (gp7*-i) is inside (**Fig. 2a; fig. S6d–l**). Outside the fingertips, the remaining residues assemble into a helical bundle comprised of 24 α-helices, with each gp7* monomer folding back on itself to contribute two helices to the bundle. There is an extensive hydrogen bond network between gp7*-o and gp7*-i through the fingertips towards the helical bundle that reinforces the scaffold of what becomes the extracellular channel (**Fig. 2b)**.

**Figure 2.**
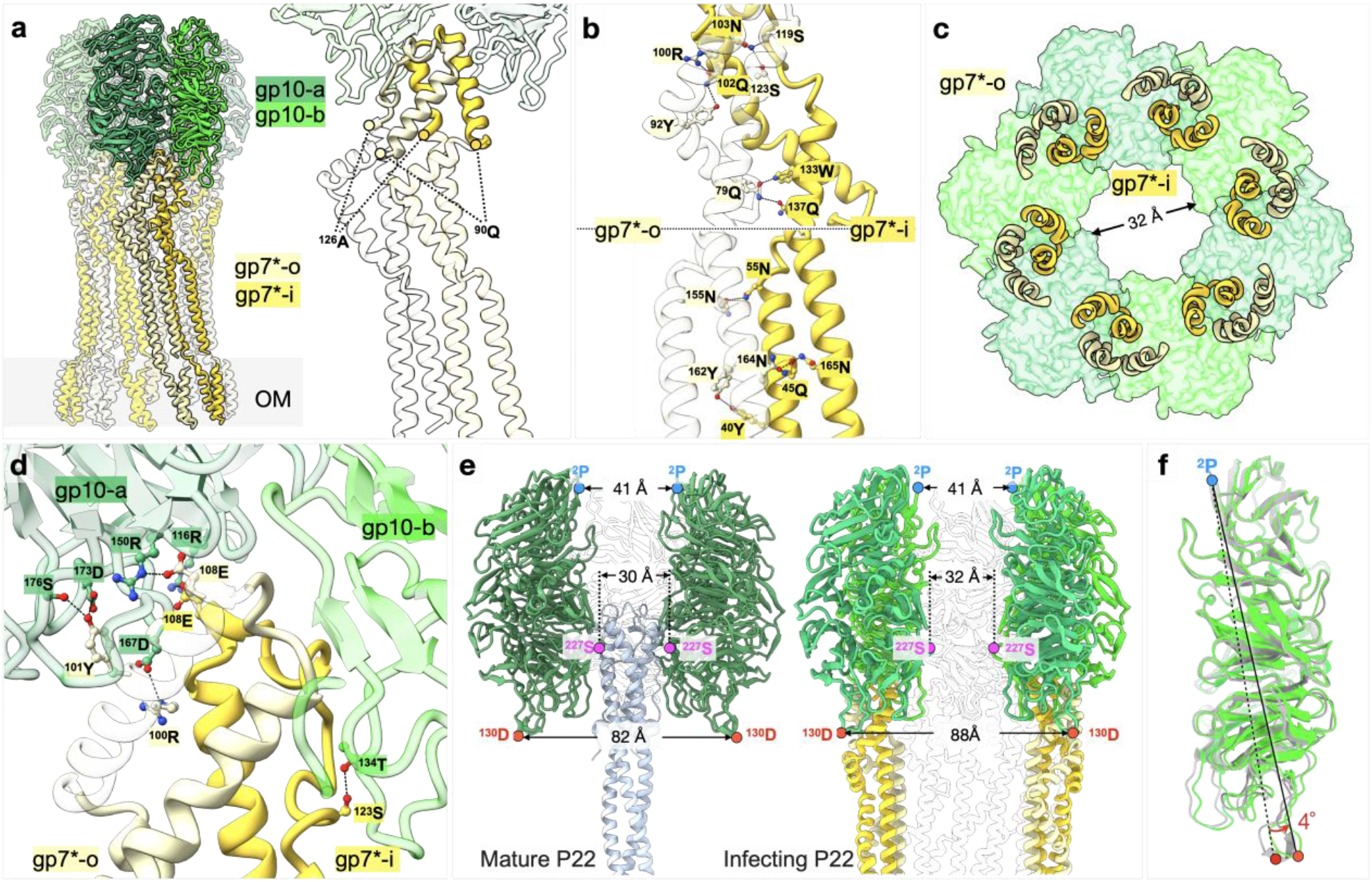
Structure of the gp10–gp7* complex. (**a**) Left: Structure of the complex formed by a gp10 hexamer (dark green and green) and a gp7* dodecamer. Adjacent gp7* molecules (gp7*-o, light yellow and gp7*-i, yellow) adopt slightly different conformations at their interface with gp10. Right: The inner and outer gp7* subunits insert into the wedge space formed two adjacent gp10 subunits. (**b**) Close-up views of gp7* show the monomers in different conformations, and highlight key H-bonds (gp7*-o - gp7*-i: ^100^R-^102^Q, ^92^Y-^102^Q, ^119^S-^103^N, ^123^S-^103^N, ^79^Q-^133^W, ^155^Q-^45^N, ^160^Q-^45^Q, ^160^Q-^165^N, ^164^N-^45^Q, ^164^N-^165^N, ^162^Y-^40^Y). (**c**) Bottom view of the interface between gp10 and gp7*, shows how gp7*-o and gp7*-i accommodate the C12–C6 symmetry mismatch with gp10. (**d**) Close-up of the gp10–gp7* interface highlights key H-bonds between gp10 and gp7* (gp10-a - gp7*o: ^176^S-^101^W, ^173^D-^101^W, ^167^D-^100^R, ^150^R-^108^E, gp10-a - gp7*-i: ^116^R-^108^E, gp10-b/gp7*-i: ^134^T-^123^S). (**e**) Conformational changes in gp10 resulted from needle release and gp7* ejection. Left: Distances between opposing gp10 subunits in mature phage. Measurements are taken at the most distal residues: ^2^P and ^130^D, and ^227^S, which makes the closest contact of gp10 with the N-terminal loop of the needle (gp10-gp26, adapted from PDB: 8EAP). Right: Similar measurements taken from infecting phage. (**f**) Superimposition of the gp10 structures from the mature (grey) and infecting (green) phages reveals a conformational change accompanied by a 4° rotation around ^2^P.

Structural plasticity of the fingertip enables extensive interactions between gp7* and gp10 (**Fig. 2c, d**). Several residues of gp7*-i interact specifically with one gp10 (gp10-a), which also interacts with two gp7*-o residues; in addition, the other gp10 (gp10-b) and gp7*-i interact **(Fig. 2d)**. Collectively, the interactions between gp10 and gp7*-o and -i stabilizes the gp10–gp7* complex and may contribute to sealing the extracellular channel, a configuration that might help prevent DNA leakage during genome translocation into the infected cell.

Conformational changes in gp10 occur before or during gp7* ejection (**Fig. 2e; fig. S3a, b**). Opposing ^2^P residues of the hexameric gp10 at the hub top remain fixed in position but the distance between opposing ^227^S residues in the center of the hub increases from 30 Å to 32 Å, and the opposing ^130^D residues at the hub base increase their separation by 6 Å to 88 Å. These changes alter the interaction of the hub with the needle from that seen in mature phage(*23*). Further, as each gp10 monomer rotates by approximately 4° outward around the ^2^P axis (**Fig. 2f**), it produces a subtle hinge-like motion that increases the hub lumen diameter, loosening its interaction with the needle. As the needle is the “plug” that confers stability to the contents of the capsid, loosening the gp10-gp26 interaction necessarily facilitates later steps in the infection pathway.

Taken together, our results define a coordinated molecular rearrangement by which P22 transforms from a pre-genome ejection to an ejection or post-ejection state, and illustrates how P22 initiates the release of all four E proteins. Structural remodeling of the tail components – through gp26 release from gp10(*23*), gp10 expansion and gp7* attachment – converts the mature virion into an infection-active state, thereby paving the way for the stepwise release of the remaining E proteins and ultimately, delivery of the genome into the infected cell cytoplasm.

### Gp20 stabilizes the gp7* outer membrane channel and creates a periplasmic channel

Gp20 was shown to span the periplasm after infection(*11*), but as a gene *16* amber mutant was used in that experiment it is unclear whether the 171 residues of the amber peptide contribute to the electron density observed. Using a mutant particle that completely lacks gp16, we can now conclude that the ejected gp20, for the first time also clearly resolved here as a dodecameric assembly, extends across the entire periplasm. Our model shows that the gp20 assembly is approximately 32 nm in length; it stabilizes the extracellular channel **(Fig. 3a)**, forms most of the channel across the outer membrane, and comprises the entire periplasmic channel **(Fig. 3b)**.

**Figure 3.**
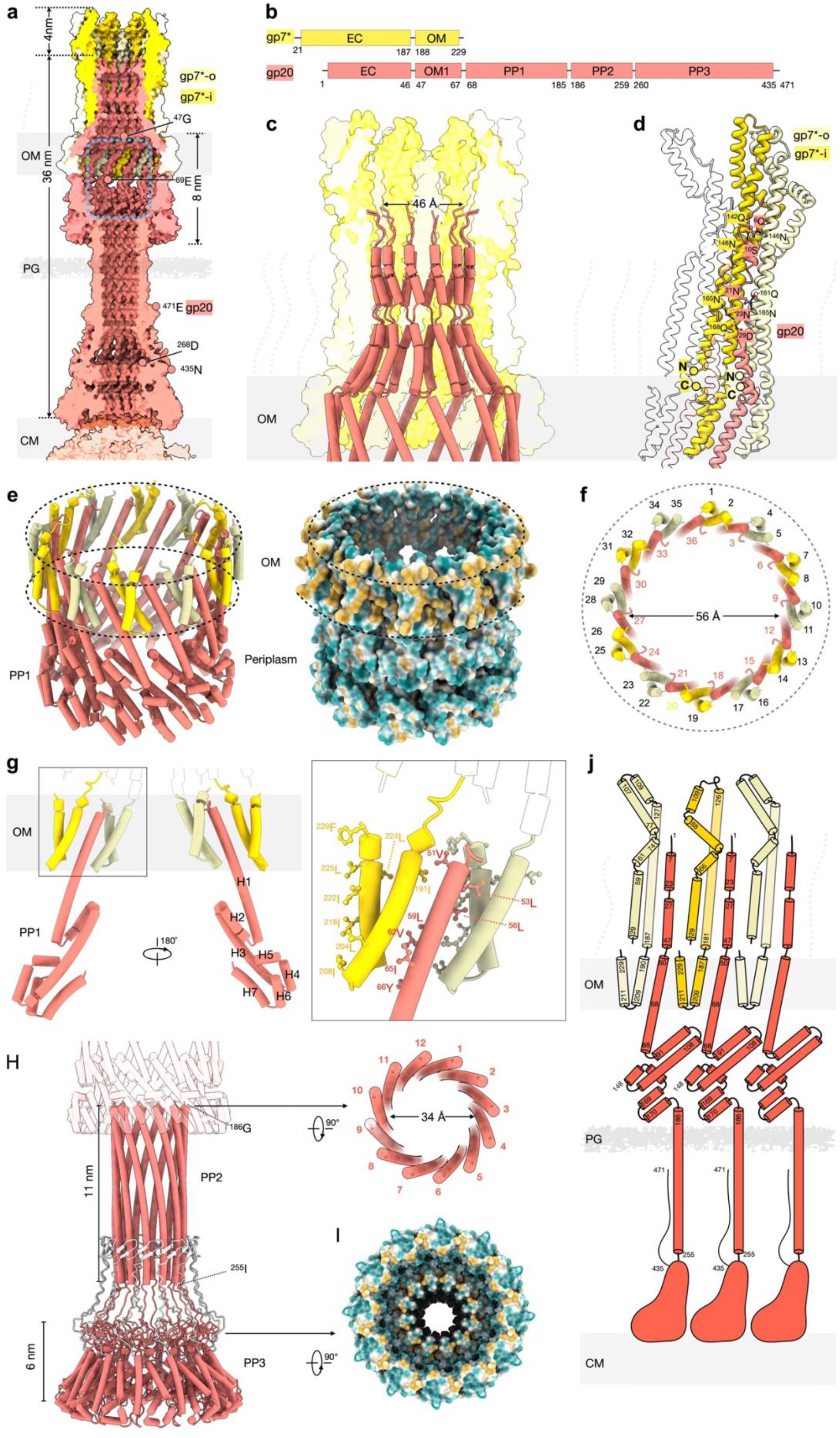
Structure of the gp7*–gp20 complex. (**a**) Medial cross-sectional view in surface rendering. gp7*-o, light yellow; gp7*-i, yellow; gp20, orange. The dotted boxed region highlights the chamber in the OM that extends into the periplasm. (**b**) Schematic domain gene product organization of gp7* and gp20. Both gp7* and gp20 contribute to extracellular (EC) and OM domains, whereas gp20 extends farther into the periplasmic space. PP1-3: periplasmic domains of gp20. (**c**) Close-up view of the EC ∼46 Å diameter channel, highlighting the internal location of gp20. Gp7* is shown in a transparent surface rendering. The N-terminal region of gp20 exhibits strict C12 symmetry and is surrounded by twelve copies of gp7*. Although gp7* assembles as a dodecamer, its extracellular domains do not adopt perfect C12 symmetry, rather the protein exhibits C6 symmetry around paired gp7*-o and gp7*-i subunits. (**d**) Close-up of the gp20 N terminus reveals key H-bonds between gp7* and gp20 in the extracellular channel (gp7*-o/gp20: ^146^N-^6^Q, ^161^Q-^21^N, ^165^N-^21^N; gp7*-i/gp20: ^142^Q-^10^S, ^146^N-^10^S, ^165^N-^23^N, ^168^Q-^29^D). (**e**) Tilted view of the pore. Hydrophobicity surface analysis illustrates the hydrophobic nature of its outer face, consistent with its outer membrane environment. (**f**) Top view of the pore showing that the transmembrane domain consists of 36 helices: 24 from 12 copies of gp7* (each subunit folds into two helices that are embedded in the OM), and 12 from gp20. (**g**) Side views of the gp7* OM domain and the gp20 OM and PP1 domains, revealing seven helices in the PP1 domain. Hydrophobic residues highlighted and labeled (gp7*: ^191^I, ^204^L, ^208^I, ^218^I, ^222^I, ^224^L, ^225^L, ^229^F; gp20: ^51^V, ^53^L, ^56^L, ^59^L, ^62^V, ^65^I, ^66^Y). (**h**) Close-up (left) and top view (right) reveal a 12-helix barrel architecture of the extended periplasmic channel formed by the gp20 PP2 and PP3 domains. PP2 spans ∼11 nm, whereas PP3 is ∼6 nm. At low confidence, AlphaFold3 predicts that the extreme C-terminal segment (residues 435– 471, light gray) folds back against the protein body (**fig. S8a**). (**i**) Hydrophobicity surface analysis of the bottom view of gp20 reveals hydrophobic residues of gp20 PP3 oriented toward the cytoplasmic membrane (CM). (**j**) Schematic representation of three gp20 with adjacent gp7*, illustrating the helix topology within the transmembrane channel.

The N-terminal 47 residues of gp20 are inserted into the lumen of the gp7* scaffold, where they form the extracellular channel (**Fig. 3c; fig. S6k, l**). However, gp20 makes no direct contact with the gp10 hub and is separated from it by ∼4 nm (**Fig. 3a**). Together with the 32-nm-long gp20 dodecamer, this results in a ∼36 nm channel extending from the tail hub to the outer face of the cytoplasmic membrane. The upper portion of the gp7*–gp20 complex is reinforced by an extensive hydrogen-bond network (**Fig. 3d**), stabilizing the extracellular channel.

The interactions between gp7* and gp20 extend deep into the outer membrane, where 36 transmembrane α-helices form what we term the outer membrane pore (**Fig. 3e, f**), which was also observed previously(*11*). Twelve of these helices are contributed by gp20, whereas the remaining 24 are formed by the two arms of the V-shaped C terminus of the dodecameric gp7*. This V-shaped configuration positions both gp7* arms within the outer membrane where they interact with adjacent gp20 helices. Eight hydrophobic residues from gp7* and seven from gp20 are embedded within the outer membrane bilayer (**Fig. 3g**), consistent with the strongly hydrophobic surface properties of this region (**Fig. 3e**).

Gp20 residues 68–185 (PP1, **Fig. 3b**) form an intertwined α-helical bundle comprising seven helices beneath the outer membrane pore (**Fig. 3g; fig. S6k)**. Together, the bundle and pore together span 8 nm along the channel axis (**Fig. 3a**), forming a chamber ∼56 Å at its widest point. Residues 186–255 (PP2, **Fig. 3b**) adopt an elongated helical barrel about 34 Å in internal diameter that crosses the peptidoglycan layer and periplasm to connect to the C-terminal plinth of gp20 (**fig. S6l**). The junction with the C-terminal plinth appears relatively dynamic (**fig. S3d–f**), suggesting that the barrel terminus may transition between two conformations, ultimately completing an open DNA translocation channel.

The C-terminal domain, residues 256–435, of gp20 (PP3, **Fig. 3b**) could not be modeled directly from the cryo-EM map due to limited local resolution. However, a high confidence AlphaFold3-predicted model fits well into the corresponding density (**fig. S6l; fig. S8b)**. AlphaFold3 predicts, albeit with low confidence, that the extreme C-terminal segment (residues 436–471) folds back against the gp20 body, a configuration that appears unlikely in the context of the cytoplasmic membrane. By contrast, AlphaFold3 predictions incorporating both the gp20 PP3 domain and the N-terminal region of gp16 (residues 1–205) suggest instead that this segment folds towards the cytoplasmic membrane and wraps around the plinth. Regardless of the precise configuration of this extreme C-terminal segment, the bulk of the C-terminal region of each gp20 monomer adopts a club-shaped structure (**Fig. 3j**) that self-associates to form a dodecameric plinth abutting the cytoplasmic membrane (**Fig. 1e, f**; **Fig. 3a**).

### *In situ* structures of E protein mutants

The periplasmic channel was resolved at 5–8 Å resolution by *in situ* single-particle cryo-EM, but the density crossing the cytoplasmic membrane was substantially less well resolved, preventing an unambiguous structural assignment (**Fig. 4a**). Further focused 3D classification using a spherical mask centered on the cytoplasmic membrane identified a distinct class, in which an additional stump-like density, resolved at ∼10 Å resolution, extends across the cytoplasmic membrane (**Fig. 4a; fig. S4b**). This density likely corresponds to gp16, which could not be detected in our earlier lower resolution study(*11*). To test this assignment, we analyzed adsorbed Δgp16 virions using the same *in situ* cryo-EM approach and applied the same 3D classification strategy. No corresponding class containing the trans-cytoplasmic membrane extension was identified, although the resulting ∼9 Å structure of the periplasmic channel closely resembles that of wild-type virions. Gp20 is therefore sufficient to assemble a stable periplasmic channel and, most importantly, gp16 alone is now shown to form the channel spanning the cytoplasmic membrane, thereby completing the complete DNA translocation channel from the phage gp10 baseplate into the infected cell cytoplasm.

**Figure 4.**
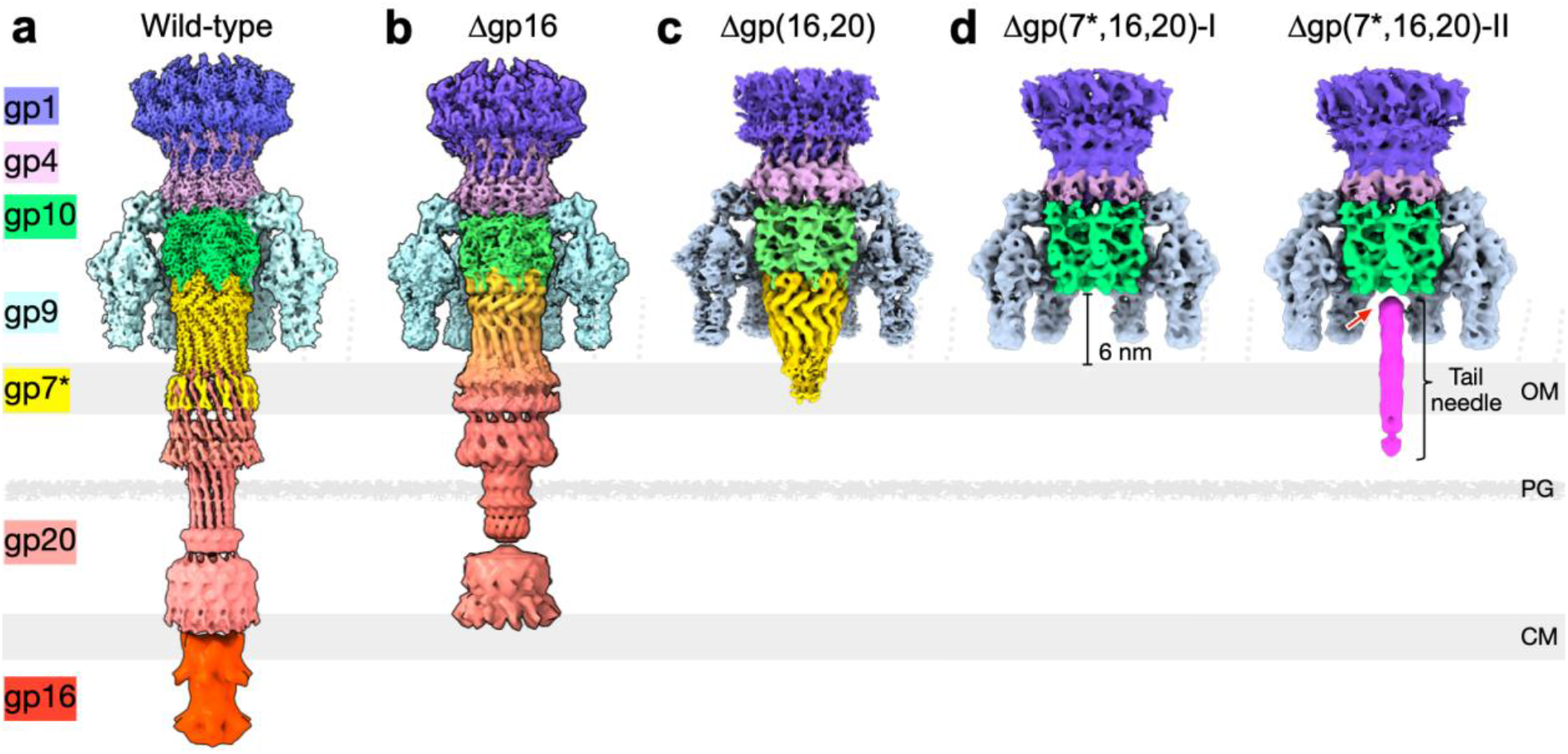
Comparative structural and functional analyses of P22 mutants. (**a**) *In situ* cryo-EM structure of wild-type P22 after gp7*, gp20 and gp16 have assembled a ∼50 nm trans-envelope channel connecting gp10 to the cytoplasm. (**b**) *In situ* cryo-EM structure of a Δgp16 infection intermediate showing a gp7*-gp20 channel. (**c**) *In situ* cryo-EM structure of an absorbed Δgp(16,20) particle in which gp7* forms an incomplete extracellular channel (arrow). (**d**) Approximately two-thirds of adsorbed Δgp(7*,16,20) particles have ejected their needle, whereas approximately one-third retain a tail needle that is uncoupled from the tail hub, resulting in a visible gap (arrow). PG, peptidoglycan.

To better understand the role of gp20 in assembly and stability of the outer membrane pore and the extracellular channel, we performed cryo-ET and subtomogram averaging on adsorbed Δgp(16,20) virions, where gp7* is the only internal E protein (**Fig. 4c**). The resulting 4.7 Å resolution structure reveals that the extracellular channel made by gp7* terminates in a cone, distinctly different from the structure seen after infection with wild-type phage where gp20 is present (**Fig. 4a**). Although only residues 35–181 of gp7* could be resolved, the secondary structure of the gp7* dodecameric complex reveals the fingertips of each gp7* dimer interacting with gp10 as in wild-type phage (**fig. S9a)**. This result strongly reinforces our conclusion that gp20 stabilizes the extracellular channel mainly formed by the gp7* dodecamer.

To determine what occurs during infection in the absence of all three internal E protein, we examined adsorbed Δgp(7*,16,20) particles by cryo-ET and subtomogram analyses. Two distinct classes were identified based on the presence or absence of the gp26 needle (**Fig. 4d**). In the major class (∼62%), neither the needle nor an extracellular channel was observed (**Fig. 4d**, left). In the minor class (∼38%), the needle was visible but was separated from the tail hub by ∼2 nm (**Fig. 4d**, right; **fig. S9b**). Compared to the mature virion, where the gp26 needle is inserted ∼39 Å into the gp10 lumen (**fig. S9c**), the needle in this intermediate has moved towards the infected cell, disengaging from the gp10 hub while its distal tip penetrates the outer membrane. Although the C-terminal segment (residues 141–233) of the needle was not resolved, most of the trimeric needle fitted well into the cryo-ET density **(fig. S9b)**, suggesting that its distal end becomes flexible or undergoes conformational distortion upon engaging the host outer membrane during this intermediate stage of needle ejection.

Collectively, our structural data strongly support the model in which the extracellular channel, outer membrane pore, and periplasmic channel are formed by just two E proteins – gp7* and gp20. They also provide a basis for a better understanding of the early process of infection initiation. To prevent premature genome leakage, assembly of the gp7* extracellular channel must be tightly coordinated with needle release and assembly of the gp20 dodecamer.

### Genome leakage associated with infecting mutant particles

The structural models we have developed for P22 infection suggest that without a full complement of internal E proteins, there is a possibility that the phage genome may escape into the extracellular medium. To test this idea directly we measured the release of P22 ^3^H-DNA during infection of cells lacking a periplasmic endonuclease (Endo I). Neither wild-type P22 nor pseudo-wild-type particles of Δ(*7,20,16*) grown on IJ612 (pTP109), where the plasmid provides gp7*, gp20, and gp16, release significant amounts of DNA into the growth medium (**Fig. 5a**). By contrast, more than 70% of phage genomes leaked from particles lacking all three internal E proteins, a result consistent with there being an ∼6 nm gap between the bottom of the gp10 hub and the outer membrane. Most particles lacking gp7* also allow their genome to leak into the growth medium; as these particles cannot make a stable extracellular channel, this result is also expected. In the absence of gp20 a limited amount of genome leakage occurs under our assay conditions, an observation supporting our conclusion that gp20 is required to complete and stabilize the gp7* extracellular channel. Particles lacking only gp16 do not release their DNA into the medium. The genome end may become trapped inside the lumen of the channel that extends from the gp10 hub to the cytoplasmic membrane or it may be released into a periplasm that in these experiments lacks Endo I.

**Figure 5.**
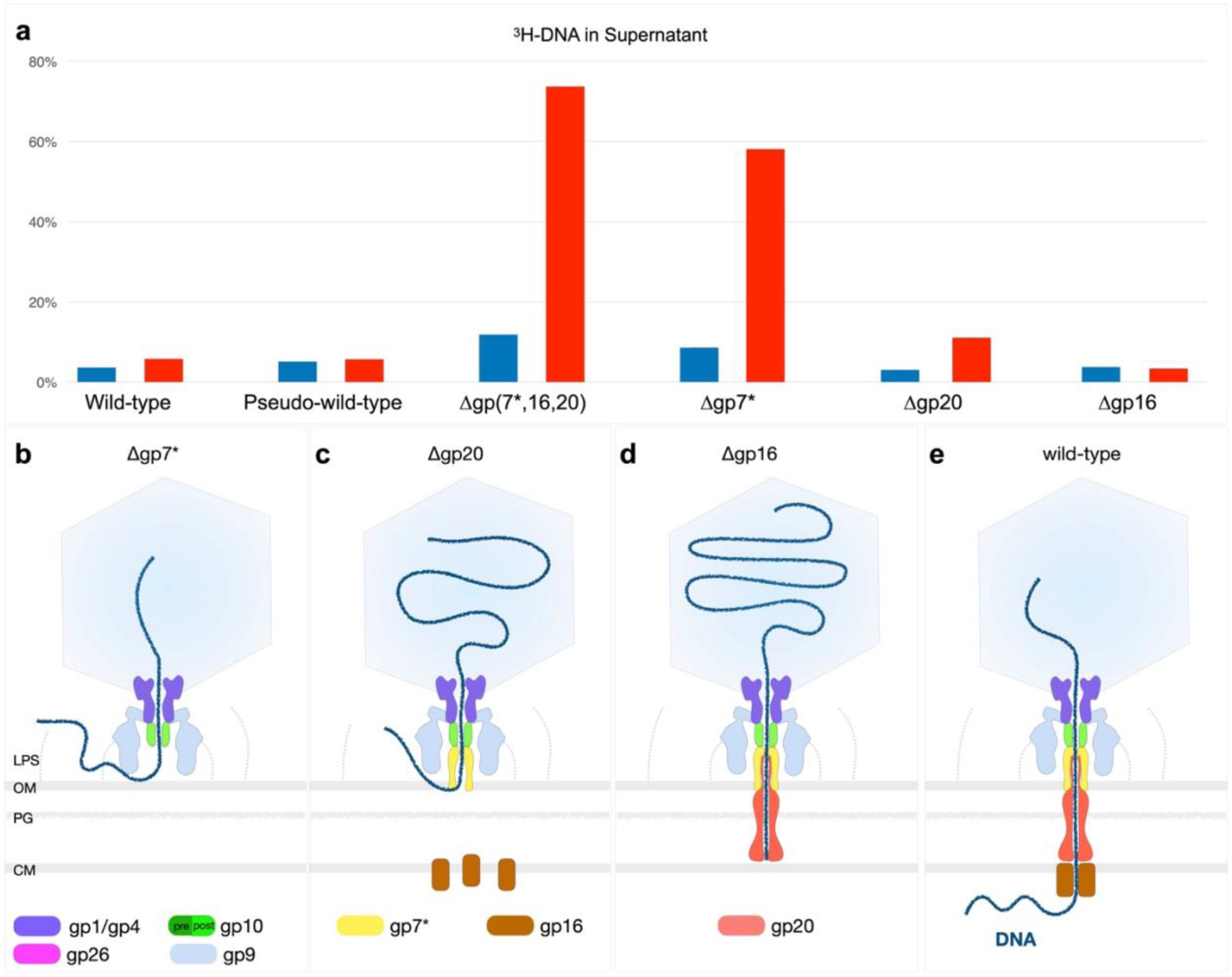
DNA release by wild-type P22 and ejection-protein mutants. (a) Phages containing ^3^H-labeled DNA were used to infect the *endA* strain TH951 at an moi of ∼4. Cells were grown to 2 × 10^8^ cells/ml at 37 °C, cooled to 10 °C immediately before infection, and treated with 100 µg/ml chloramphenicol to prevent phage development. Following adsorption for 10 min, samples were collected and the remaining infected cells were returned to 37 °C and incubated for 60 min. Blue: % DNA in supernatant immediately before shifting back to 37 °C; red: % DNA after 60 min infection at 37 °C. Adsorption and genome leakage model for Δgp7* (**b**), Δgp20 (**c**), Δgp16 (**d**), and wild-type (**e**) virions.

### Extra-cytoplasmic complementation of P22 mutants

Growth of P22 mutants defective in any of the genes *7, 20*, or *16* result in morphologically normal particles that are, however, non-viable. Decades ago, it was shown that under non-permissive conditions an infecting P22 gene *16*(Ts) or gene *16* amber mutant was complemented in *trans* by a co-infecting P22 gene *20* amber mutant(*27, 28*). Complementation was shown to occur extra-cytoplasmically by the defective phages, prior to genome internalization in the infected cell. Complementation therefore occurred before any phage gene expression in the infected cell. The converse reaction: complementation of a P22 *20* mutant by a confecting P22 *16* amber mutant did not occur. Thus, only gp16 can act *in trans*. Particles lacking gp7* have been said(*20, 35*) not to complement, or be complemented by, other mutants but no data have been presented. More importantly, no molecular explanation for any of these results has yet been offered.

Although particles containing gp16(Ts) do not support phage growth under non-permissive conditions, it is possible that they retain partial activity. Similarly, it is possible that the gene *16amN121* amber peptide (171 residues) is packaged into virions and retains some biological function. To exclude these possibilities, we performed a series of complementation tests using particles formed in the complete absence of one or more E protein (**Table 1a**). Although several independent preparations of Δgp7* particles yielded higher apparent yields of progeny than expected, coinfections of Δgp7* with either Δgp20 or Δgp16 yielded no additional progeny. In other words, Δgp7* particles do not complement, and are not complemented by, coinfection with defective particles lacking a different E protein. A very different result is obtained, however, after coinfection with Δgp20 and Δgp16, when a substantial burst of progeny phage is released. This result is in full agreement with earlier experiments(*27, 28*), where it was concluded that gene *20*- and gene *16*-defective particles complement in coinfections. However, our experiments do not allow us to identify which phage or phages are present among the progeny because the parent genome is the same in both input phages (different plasmids are used to make the different defective phage particles whose genomes lack all three internal E protein genes). We therefore did additional complementation experiments using amber mutant particles (**Table 1b**). These data are comparable to previous experiments for the combinations of gene *20-* and gene *16*-defective particles(*27, 28, 35*) but extend their data by showing that neither available gene *7* amber mutant can successfully participate – either as donor or recipient – in complementation experiments and by excluding the possibility that the amber peptides of gp20 and gp16 have any critical function in facilitating complementation.

**Table 1.** Extra-cytoplasmic complementation.

| **Phage Particle** | **Progeny Phage/Cell** |  |
| --- | --- | --- |
| **(a)** | **Test 1** | **Test 2** |
| P22<sup>+</sup> | 228 | 260 |
| Δgp7* | 25 | 15 |
| Δgp16 | 4 | 13 |
| Δgp20 | 4 | 2 |
| Δgp7*+Δgp16 | 29 | 14 |
| Δgp7*+Δgp20 | 21 | 24 |
| Δgp16+Δgp20 | 79 | 94 |
| **(b)** |  |  |
| Wild-type | 171 | 209 |
| *7amH1363* (gp7<sup>-</sup>) | 1 |  |
| *7amH1365* (gp7<sup>-</sup>) | 8 |  |
| *20amN20* (gp20<sup>-</sup>) | <1 | <1 |
| *16amN121* (gp16<sup>-</sup>) | <1 | <1 |
| *7amH1363* + *16amN121* | 5 |  |
| *7amH1365* + *16amN121* | 15 |  |
| *7amH1363* + *20amN20* | 2 |  |
| *7amH1365* + *20amN20* | 11 |  |
| *16amN121* + *20amN20* | 116* | 134* |

Experiments used an estimated multiplicity of 20 for single infections and 10 for each phage in coinfections. Independent phage stocks were used in tests 1 and 2.
<sup>a</sup> All phages carry the *c1-7*, *13amH101*; those containing the prefix Δ also contain the Δ(*7,20,16,sieA*) mutation, with the deletion being partially complemented by plasmid-borne genes to give particles lacking only one E protein (see Materials and Methods)
<sup>b</sup> All phages carry the *c1-7* and *13amH101* mutations plus the E gene mutation indicated. The *7amH1363* amber mutation is at codon 78, that in *7amH1365* is at codon 155; the *20amN20* mutation affects codon 79 and *16amN121* codon 172.
\* 92 of 92 progeny phages in the mixed infections were shown by a subsequent standard complementation test using the Su<sup>-</sup> host IJ612 and mutant P22 strains to be exclusively *16amN121*.

## Discussion

Our innovative *in situ* cryo-EM approach enabled visualization of P22 infection intermediates at unprecedented resolution, providing the structural basis for a mechanistic model of infection initiation (**fig. S10; Movie S2)**. Following phage adsorption and ejection of internal E proteins, the portal barrel collapses, providing direct structural evidence that the portal and tail lumen serves as a storage site for the internal E proteins(*23*). Analyses of E protein mutant virions reinforces this conclusion. Importantly, our near-atomic structures reveal a conformational change in the gp10 hexamer that triggers infection by allowing release of the gp26 needle. Needle release then allows ejection of gp7* and its assembly into a dodecameric channel bridging the gp10 tail hub and the bacterial outer membrane.

These observations indicate that gp26 release and gp7* channel assembly are tightly coordinated during infection. In virions lacking all three internal E proteins, we observed a substantial population where the needle was clearly separated from the gp10 hub but remained embedded in the outer membrane. Although not yet visualized using wild-type phage, this structure must exist, if only fleetingly, during the normal ejection of gp26 into the infected cell periplasm. This suggests that efficient and complete translocation of gp26 into the periplasm may require the ejection and assembly of gp7* into the extracellular channel. In the absence of any internal protein that can be ejected, genome ejection could theoretically facilitate entry of the complete needle into the periplasm but as DNA has no intrinsic propensity to penetrate a membrane it may simply diffuse away from the phage-cell complex. In this situation there may therefore be a lesser incentive for the needle to complete its entry into the periplasm, allowing it to be visualized as a distinct intermediate structure.

We propose that, as the virion reorients on the cell surface from its initial obliquely bound state on the cell surface to the perpendicular, the tail needle is forced into the outer membrane (**fig. S10**). Although the distal end of the needle may be somewhat flexible(*36, 37*), continued reorientation of the capsid will eventually cause the proximal end of the tail needle to exert a mechanical force on the hexameric gp10 hub. A likely result of this force is the observed ∼4° outward shift of gp10 monomers, which will weaken the plug function of gp26 and trigger its ejection. This is rapidly followed by ejection of gp7* from the portal lumen and its oligomerization into the extracellular channel.

The gp7* dodecamer alone forms the extracellular channel and although it penetrates the outer membrane its stability in that location requires the N-terminal region of the dodecameric gp20. The inner leaflet of the outer membrane is deformed by gp7* and gp20, which together form a chamber (in cross-sectional view, a membrane pore) whose function, if any, can only be speculated. The chamber is followed by a sudden narrowing into the helical barrel region consisting of only gp20. The dodecameric gp20 forms a channel across the entire periplasm, extending to the outer face of the cytoplasmic membrane. The inner diameter of the channel is more than sufficient to allow the passage of a dsDNA helix. Because we see no cytoplasmic membrane penetration by gp20 we hypothesize that its “club foot” (**Fig. 3**) serves as a template for assembly of a gp16 complex that completes the P22 DNA translocation channel from the gp10 baseplate into the infected cell cytoplasm. AlphaFold predictions of combinations of 12 copies of the gp20 PP2–PP3 domains with 6 copies of gp16 suggest that they all form a continuous channel-like structure extending from gp20, with an inner lumen suitable for DNA passage through the cytoplasmic membrane **(fig. S8)**.

Sf6 is a close homolog of P22, and the counterpart protein to P22 gp20 in Sf6 is gp12; the two proteins exhibit 46% amino acid sequence identity over their N-terminal halves although their C-termini are less similar (**fig. S11**). We have demonstrated that the N terminus of dodecameric P22 gp20 interacts intimately with dodecameric gp7*, whose counterpart is Sf6 gp11; the latter is 82% identical by sequence to P22 gp7 and thus likely forms an equivalent dodecameric extracellular channel. These data strongly suggest that the periplasmic channels made by P22 gp20 and Sf6 gp12 are structurally similar and are both dodecamers. However, Sf6 gp12 was purified using a plasmid expression vector and shown to assemble as a decamer that features a tube and terminal crown domain(*38*). The overall length of this structure is only 15 nm, far too short to completely span the periplasm as its N terminus would be expected to interact with Sf6 gp11 in the OM. Thus, assembling Sf6 gp12 *in vitro* probably does not yield a biologically active form.

### A unique extra-cytoplasmic complementation experiment explained by structure

In a normal productive infection, a wild-type P22 genome does not become exposed to extra-cytoplasmic nucleases. Only when genome translocation into the cytoplasm is blocked, for example during P22 superinfection of a P22 *sieA^+^* lysogen or T4 superinfection of a previously T4-infected cell does the ubiquitous (in enteric bacteria) periplasmic Endo I have a possibility of attacking the superinfecting genome. The tail tube of myophages like T4 may directly protect the genome from periplasmic nucleases, but podophages like P22 have to assemble a trans-periplasmic channel *de novo*. This channel not only protects the genome from Endo I but also provides a direct route from the body of the phage into the infected cell cytoplasm.

The channel made by P22 is comprised of three E proteins that in mature phage are primarily stored in the portal barrel and the portal-tail lumen. In the absence of any E protein, there is a ∼6 nm space between the bottom of the gp10 hub and the cell surface through which the phage genome can escape into the extracellular media (**Fig. 4d**). In the absence of gp7*, a comparably large fraction of the infecting genomes is also released from the infected cell complex. Although gp20 and gp16 are ejected from the capsid before the DNA(*23, 39*), if they cannot interact with the gp10 hub and form an extracellular channel they presumably will also be found in the growth medium. This would result in the same gap between gp10 and the cell surface as in the absence of all E proteins, a situation again allowing the genome to leak into the surrounding fluids. The incomplete release of DNA into the media in both these experiments is not understood but some genomes may become entangled in the milieu of tailspikes and O antigen-LPS chains.

When the infecting phage contains gp7*, genome leakage into the media is largely suppressed and is reduced almost to wild-type levels when gp20 is also ejected. These results are consistent with our structural studies showing that gp7* forms much of the extracellular channel but requires gp20 to stabilize the channel in the outer membrane and to extend a channel across the periplasm. Also as expected from the structural data, the presence or absence of gp16 has no significant effect on the leakage of an infecting genome into the surrounding fluid.

More than 50 years ago, it was shown that two different defective P22 virions could complement and allow a productive infection(*27, 28*). Neither phage alone was capable of getting its genome into the cytoplasm of the infected cell, and it was clearly shown that complementation did not require new protein synthesis. We refer to this novel process as extra-cytoplasmic complementation. It was also shown that complementation between gene *16* and gene *20* mutants was unidirectional: gp16 from the gene *20* mutant virion could rescue the gene *16* mutant but gp20 was unable to work *in trans*. We can now completely explain these observations at a structural level and extend them to gp7*, which cannot participate either as a donor or recipient in extra-cytoplasmic complementation.

Gene *7* mutants cannot provide gp20 or gp16 to a coinfecting phage; those proteins are unable to enter the infected cell because there is no extracellular channel to guide them after they are ejected past the gp10 hub of the gene *7* mutant (**Fig. 5b**). Similarly, even if a gene *20* or gene *16* mutant particle ejects more gp7* than it needs to make its own extracellular channel, wherever the extra protein goes to in the infected cell, it does not have a path to exit the cell and engage with the gp10 hub of a coinfecting gene *7* mutant (**Fig. 5c, d**). Again similarly, after ejection from a capsid gp20 must immediately associate with the gp7* extracellular channel in order to stabilize it and to extend a channel to the cytoplasmic membrane. It is therefore not free to diffuse and complement another infecting particle. Furthermore, even if more than 12 copies of gp20 are ejected from one virion the extra copies must diffuse back from the periplasm into the outer leaflet of the outer membrane to complex with gp7* of another infecting particle. These theoretical possibilities seem highly improbable at best. By contrast, our experimentally determined structures provide a logical framework explain why gp7* and gp20 of an infecting particle only work *in cis*. A schematic animation of extra-cytoplasmic complementation as we now understand it is shown in **Movie S3**.

The biological observation that gp16 from one phage virion can complement a coinfecting gene *16* mutant outside the cytoplasm of the infected cell requires that the protein be free to diffuse. For this to happen in our experimental design, gp16 must be efficiently ejected into a host cell in the absence of any channel formed by gp20. This means that gp16 must pass through the nascent gp7* extracellular channel, in the absence of its stabilization and extension by gp20, and enter the periplasm where it probably diffuses to the cytoplasmic membrane. By contrast, association of the outer membrane portion of the gp7* channel with the N-terminal domain of gp20 will anchor it, allowing assembly of the periplasmic portion of the DNA translocation channel but preventing diffusion to other parts of the cell.

Gp16 from a phage containing all E proteins still complements a gp16-deficient mutant(*27, 28*), implying that gp16 is slow to associate with the gp20 periplasmic channel, more copies of gp16 are ejected than are required for infection by one particle, or gp16 is ejected before gp20. Our data using Δgp20 particles allow favoring the last possibility, although we cannot formally exclude the others. The concentration of gp16 inside proheads assembled *in vitro* appears to be fixed as providing excess gp16 does not increase its abundance(*40*). The copy number of gp16 per virion has been estimated to be ∼6-12(*41, 42*) but the composition of the cytoplasmic membrane channel, which contains the ejected gp16, has yet to be conclusively determined.

When the gp16-acccepting phage is present at higher concentrations than the gp16 donor, complementation efficiency declines(*28*), suggesting that monomeric gp16 interacts with the gp20 channel. Indeed, using disrupted gp20-minus proheads as a source, gp16 was purified and shown to be monomeric(*40*). Interestingly, this study also stated that gp16 could not be purified away from gp20 if the latter was also present in proheads. There may therefore be a mechanism, spatial or structural, to prevent the two proteins from complexing in the virion prior to genome ejection. It seems likely that gp7* should also be sequestered from gp20 while inside the capsid. How such sequestration is manifest and how the three E proteins in proheads eventually migrate to the portal lumen so that they can be ejected prior to the phage genome(*11, 39*) is not known.

Although it lacks any strongly predicted transmembrane helix, gp16 was shown to associate with a cytoplasmic membrane preparation *in vitro*(*43*). We can therefore imagine that after ejection from an infecting P22, gp16 monomers rapidly diffuse in or on the surface of the cytoplasmic membrane. It is clear that gp16 is not capable of forming an open channel across the membrane in the absence of gp20 because infection by P22 particles lacking gp20 does not affect cellular growth rate and does not lead to cell killing. Tailspike binding to O antigen is thought to be normally irreversible(*44*), making it highly improbable that an adsorbed phage that has ejected gp7* and gp20 will move significantly from its original site of adsorption. At least 700 P22 particles were estimated to bind to a single *S.* Typhimurium cell(*45*); at more common multiplicities of infection, the coinfecting phage that provides gp16 to another infecting particle is thus unlikely to be close by on the cell surface. Diffusion of gp16 must therefore be very rapid and its association constant to a periplasmic channel, or perhaps to the *Salmonella* protein YajC, must be very high in order for complementation to be successful. Interestingly, a high association constant to gp20 favors our idea that gp16 enters the cell before the dodecameric gp20 channel is assembled. YajC is a cytoplasmic membrane protein that is essential for a P22 genome to reach the cell cytoplasm(*46*). Though non-essential, YajC has been shown to be part of the Sec secretion system(*47, 48*) and is therefore expected to be a moderately abundant protein. However, it is not known if YajC directly facilitates the transport of P22 across the cytoplasmic membrane or whether its absence affects the secretion or presence of some other protein important for P22 infection.

## Supporting information

Movie S1. Animation showing in situ structures derived from cryo-ET/subtomogram averaging and in situ single-particle cryo-EM.

Movie S2. Animation showing wild-type P22 infection initiation.

Movie S3. Animation showing extra-cytoplasmic complementation by coinfecting delgp20 and delgp16. For simplicity, only six copies of gp16 are modeled.

## Acknowledgments

We thank Sherwood Casjens, not only for kindly providing phage strains but also for his unpublished sequencing data defining the amber mutations used and reported here. We thank Jennifer Aronson for critical reading of the manuscript and Shenping Wu for suggestions for cryo-EM data collection. We also thank Hang Zhao and Jack Botting for assistance with data collection and animation. Cryo-EM data were collected at Yale Cryo-EM Resources funded in part by NIH grant 1S10OD023603-01A1.

## Author contributions

Conceptualization: HY, JL, IJM Methodology: HY, MC, CW, JL, IJM

Investigation: HY, JY, TP, CW, MC, JL, IJM Funding acquisition: JL, IJM

Project administration: JL, IJM Supervision: JL, IJM

Writing – original draft: HY, JL, IJM Writing – review & editing: HY, JL, IJM

## Data, code, and materials availability

Wild-type P22 infection cryo-EM maps and atomic coordinates have been deposited in the Electron Microscopy Data Bank (EMDB) and Protein Data Bank (PDB). The locally refined extracellular channel map and coordinates containing gp1–gp4–gp9–gp10–gp7*–gp20 is available under accession codes EMD-77019 and PDB-13EF. The outer membrane and periplasmic channel map and coordinates containing gp7*– gp20 is available under accession codes EMD-76290 and PDB-12BO. The periplasmic and cytoplasmic channel map containing gp20–gp16 has been deposited in EMDB under accession code EMD-78681. The Δgp16 P22 infection overall channel cryo-EM map has been deposited in EMDB under accession code EMD-77764. The wild-type P22 infection extracellular complex and periplasmic channel cryo-ET maps have been deposited under accession codes EMD-77715 and EMD-77716, respectively. The Δgp(16,20) P22 infection extracellular complex cryo-ET map has been deposited under accession code EMD-77720. The Δgp(7*,16,20) P22 infection extracellular complexes with and without the tail needle have been deposited under accession codes EMD-77721 and EMD-77752, respectively.

## Materials and Methods

### Bacteria

IJ612 (originally MS1868): *Salmonella enterica* sv. Typhimurium LT2 *leuA414*(Am) *hsdR* (Fels2^-^) and its isogenic *supE* derivative IJ613 (MS1883) are from the lab collection. TH951 (LT2 s*erA4 endA41*::Tn*10*) was a gift from Kelly Hughes. Bacteria were grown aerobically at 37 °C in rich media (per liter: 10 g Bactotryptone, 5 g Difco yeast extract, 5 g NaCl), supplemented with antibiotics as necessary.

### Phage preparation and purification

P22 *c*I-7 *13amH101*, a clear mutant of P22 that harbors an amber mutation in the holin (lysis) gene, is referred to here as wild-type (WT) P22 or P22^+^ as no structural genes are altered. It is the parent to all other phage mutants used. P22 amber mutants were kind gifts from S. R. Casjens. Defective amber mutant particles were made by a single round of phage growth in IJ612. Other P22 strains were constructed and propagated as described previously(*23*). P22 Δgp(7*,20,16), a phage deleted for all three internal E protein genes, was propagated on IJ612 (pTP109), where the plasmid supplies the essential E proteins. This phage genome also lacks *sieA*, which encodes a protein that is not a structural component and is non-essential for phage growth; *sieA* is not pertinent to the experiments described here. Particles lacking all E proteins [P22 Δgp(7*,20,16)] were prepared by a single round of phage growth on IJ612, whereas particles lacking just one of the E proteins used as host IJ612 containing a plasmid(*23*) that provides the other two: pTP29 allows production of P22 Δgp7*, pTP68 of P22 Δgp20, and pTP28 of P22 Δgp16. Clarified lysates were treated with DNase I to reduce viscosity and phages were purified on an equilibrium CsCl-density gradient; they were stored in CsCl until just prior to use. The specific infectivity of WT P22 was determined as ∼4 × 10^11^ plaque-forming particles per OD_260_, and this value was used to estimate the concentration of non-infective mutant phage particles. ^3^H-labeled P22 particles were prepared by infecting IJ612 growing in M9 medium containing 100 µg/ml deoxyadenosine and ^3^H-thymidine (2 Ci/mmol, 10 µCi/ml; Perkin-Elmer).

### Complementation experiments

Extra-cytoplasmic complementation experiments using defective P22 particles completely lacking an E protein used the permissive host IJ612(pTP109). The data shown used an estimated multiplicity of infection (moi) of 10 of each phage (20 for WT by itself), although lower particle inputs were also successfully tested. Cells grown at 37 °C to a density of 2 × 10^8^/ml were infected and incubated for 2 hr, vortexed in the presence of CHCl_3_, and clarified by centrifugation. The supernatant was then titered using IJ612(pTP109). Extra-cytoplasmic complementation experiments using amber mutant particles were performed similarly but using the permissive host IJ613; progeny plaques were subsequently scored for their genotype by additional routine complementation tests in the non-permissive host IJ612.

### Leakage of P22 DNA from infected cells

TH951 growing at 37 °C and at a density of 2 × 10^8^ cells/ml was cooled to 15 °C, treated with 100 µg/ml chloramphenicol, and infected with WT or mutant P22 using an moi of ∼4. After 10 min to allow for adsorption, cultures were returned to 37 °C for 60 min incubation. Samples were removed before and after incubation and centrifuged rapidly to separate infected cells and supernatant fluid. Radioactivity in both fractions were quantified using a scintillation counter.

### Preparation of minicells infected by P22

IJ2299 (originally TH16943: *S*. Typhimurium LT2 *araBAD1091*::*ftsZ*^+^) was grown at 37 °C in 50 ml LB medium to generate minicells, as previously described(*11*). In brief, a 50 ml overnight culture was centrifuged twice at 3,000 × g for 5 min to remove regular cells. The supernatant was centrifuged again at 20,000 × g for 20 min; the minicell pellet was then resuspended in LB medium containing 500 μg/ml rifampicin to give an OD_600_ of 0.3–0.4 and incubated at 37 °C for up to 60 min with phages at various concentrations based on their original titer or estimated titer. 5 µL samples were then mixed with 10 nm colloidal gold and deposited onto a freshly glow-discharged grid (Quantifoil Cu 2/1 300 mesh). The grid was blotted for 8 sec before rapid freezing in liquid ethane using a homemade plunger.

### Cryo-ET data collection

Cryo-ET data were collected from minicells infected with P22 particles using a Titan Krios G2 microscope (Thermo Fisher Scientific) equipped with a field emission gun, a K3 Summit direct detector (Gatan), and a GIF energy filter (Gatan) with a 20 eV slit width. Tilt series were acquired using SerialEM(*30*) and FastTomo(*49*) in a super-resolution dose-fraction mode at a nominal magnification of ×42,000, corresponding to a calibrated pixel size of 1.074 Å. Each tilt series covered an angular range of -48° to +48° with 3° increments (33 images total) and a cumulative electron dose of ∼70 e^-^/Å^2^. Defocus values ranged from -3 and -5 μm. In total, 731, 468, and 561 tilt series were collected from minicells infected with wild-type, Δgp(16,20), and Δgp(7*,16,20) phages, respectively. An moi of approximately 30 was used for all infections.

### Cryo-ET data processing and subtomogram averaging

Movie stacks of tilt series were drift-corrected using MotionCor2(*50*) and at a binning factor of 2 to generate micrographs with a pixel size of 2.148 Å/pixel. Tilt series alignment and tomogram reconstruction were done with an automated pipeline in EMAN2(*29*). With the geometry of the tilt determined from the tilt series alignment, a contrast transfer function (CTF) was estimated for each micrograph. Locations of 37,841 phage particles were selected using ConvNet-based autopicking from tomograms, and raw subtomograms were generated from tilt series after per-particle-per-tilt CTF correction. The initial subtomogram average of the capsid was reconstructed with icosahedral symmetry. We then performed two rounds of symmetry expansion and iterative classification. The first round relaxed the symmetry to C5 to identify the unique vertex where the portal complex and tail are located. The second round focused on the portal complex and relaxed the C5 symmetry to C1, resolving the symmetry mismatch between the capsid and portal. Non-portal vertices were then extracted and refined with C5 symmetry, yielding a reconstruction at 4.3 Å resolution. Another round of subtomogram refinement was then performed on the entire particle with C6 symmetry after shifting the box center to the phage tail, using the symmetry-relaxed C1 structure and orientation as a starting reference. Multiple rounds of multi-reference refinement resolved two distinct conformations of the tail: with and without the outer membrane. The membrane-associated class was further refined to a subtomogram average at 8.7 Å resolution by gathering sub-tilt alignment information derived from the non-portal vertex refinement. A similar strategy was applied after shifting the particle box center to the periplasmic channel, yielding a reconstruction of the complete channel at ∼16 Å resolution. The processing workflow and resolution determination summary is shown in **fig. S1a–d**. The Δgp(16,20) dataset was processed using EMAN2; 12,650 particles were auto-picked and processed to yield a 4.7 Å reconstruction of the extracellular complex (**fig. S1e)**. The Δgp(7*,16,20) data were also processed using EMAN2, with 38,750 particles auto-picked. Multi-reference refinement focused on the tail region and resolved both needle-containing and needle-lacking classes. Statistics of cryo-ET data collection and processing are summarized in **Table S1.**

### Cryo-EM sample preparation and data collection

For cryo-EM, 5 µL of phage-minicell suspension was applied to a glow-discharged grid (Quantifoil Cu 2/1 300 mesh). The grid was blotted for 6 sec under 90% relative humidity and plunge-frozen in liquid ethane using a GP2 automated freezer (Leica). Cryo-EM samples were initially screened on a 200 kV Glacios microscope (Thermo Fisher Scientific) for optimal ice thickness and minicell concentration. Typically, an moi of approximately 30 was optimal for ensuring that phage particles adsorbed to minicells would be readily visible in the thinnest regions of vitreous ice. Grids meeting these criteria were subsequently transferred to a 300 kV Titan Krios G2 microscope for high-resolution cryo-EM data acquisition.

Conventional single-particle cryo-EM data collection typically acquires images automatically from predefined acquisition points within regularly patterned holes. For sparsely distributed targets such as phage-infected minicells, however, this strategy results in most images lacking targets of interest. To overcome this limitation, data were acquired using SerialEM(*30*) together with a customized multishot script(*31*), in which acquisition targets were manually selected at minicell edges bearing visibly adsorbed phage particles in thin vitreous ice. This targeted acquisition strategy substantially increased data-collection efficiency, yielding approximately 6,000 usable micrographs per day. Images were recorded with a defocus range of -1.6 to -2.2 μm and a total electron dose of 55 e⁻/Å² per movie. Each movie was collected over 2.6 sec and fractionated into 40 frames (0.065 sec per frame). For wild-type P22-infected minicells, two datasets comprising 4,555 and 8,743 movie stacks were collected at a nominal magnification of ×81,000 (calibrated pixel size 0.534 Å) and ×64,000 (calibrated pixel size 0.673 Å), respectively. For Δgp16-infected minicells, 10,453 movie stacks were collected for at a nominal magnification of ×81,000, corresponding to a calibrated pixel size of 0.534 Å.

### Cryo-EM data processing

Motion correction was performed in cryoSPARC v4.6.0(*34*) with 2× binning, resulting in a pixel size of 1.068 Å/pixel for data processing, followed by CTF estimation using Patch CTF Estimation in cryoSPARC (**fig. S2)**. A manually curated subset of 546 capsid particles underwent 2D classification to generate templates for Topaz(*51*) training, enabling automated picking of 322,345 particles via Topaz Extract. Motion-corrected micrographs and selected particles were imported into RELION 4.0.1(*33*) via pyem(*52*). Tail-orientation determination was performed as described(*53*). In brief, particles were re-extracted in RELION from motion-corrected micrographs imported from cryoSPARC, after which CTF estimation was performed with CTFFIND4(*54*), using the imported particle coordinates. An initial model was generated using icosahedral symmetry (I3), followed by 3D auto-refinement. At this step the tail orientation is distributed among all capsid vertices. Reconstructed particles obtained after 3D refinement were symmetry-expanded, resulting in a 60-fold increase in particle number relative to the original dataset. A spherical mask centered on a single vertex was applied in EMAN2 to focus tail signal convergence. The symmetry-expanded particles were then subjected to non-image-aligned 3D classification, with the regularization parameter T and the number of classes both set to 10. Only classes containing the portal-tail complex were selected for subsequent processing. After removing duplicates, this procedure yielded 49,494 particles with a localized capsid-tail vertex that were subsequently imported into cryoSPARC. Following homogeneous reconstruction, C5 symmetry expansion and 3D classification were applied to identify tails with C6 symmetry. Extended-tail particles from 3D classes with different rotational orientations were then aligned using Align 3D Maps in cryoSPARC. Iterative local refinement and local CTF refinement under C6 symmetry resolved the extracellular channel of the gp9–gp10–gp7*–gp20 complex at 3.10 Å resolution with 685,062 symmetry-expanded particles. The structure of the portal-adaptor complex was further optimized under C12 symmetry, achieving a final resolution of 3.15 Å.

The volume alignment tool re-centered particle boxes to focus on the OM region. Surrounding membrane bilayers often obscure transmembrane protein density in native-state cryo-EM reconstructions, and this effect is especially pronounced for phage-generated channels, which, as foreign protein assemblies are likely to be less stably embedded in the bacterial envelope than endogenous proteins. To reduce membrane-derived interference, we performed membrane-signal subtraction before further analysis. An OUTER MEMBRANE mask was segmented from the homogeneous reconstructed map using ChimeraX(*55*) and Volume Tools in cryoSPARC(*34*), and was subsequently applied to subtract the OUTER MEMBRANE density and improve the signal-to-noise ratio of the transmembrane protein density. Subsequent symmetry expansion and 3D classification removed poorly aligned particles, enabling local refinement of the OUTER MEMBRANE and periplasmic channel of the gp7*–gp20 complex at 3.99 Å resolution using 519,368 symmetry-expanded particles. The particle box was then recentered to the periplasmic space, and symmetry expansion and 3D classification removed incomplete channel particles, enabling local refinement of the OUTER MEMBRANE and gp20 periplasmic channel, including PP1 and PP2, to 5.22 Å resolution from 120,875 symmetry-expanded particles. This proved sufficient to resolve secondary-structure features. Subsequent 3D classification, focused on the entire periplasmic channel, resolved three distinct conformational classes of the periplasmic channel (**fig. S5d–f**). The most complete class was locally refined to an overall resolution of 8.10 Å. Following C12 symmetry expansion, focused 3D classification using a spherical mask centered on the cytoplasmic membrane identified a class containing an additional density spanning the cytoplasmic membrane (**fig. S4b**). Detailed cryo-EM processing and resolution-estimation workflows are shown in **fig. S2,3**. Local resolution distributions were estimated in cryoSPARC and are shown in **fig. S5,6**. Using the same strategy, 87,697 capsid particles were autopicked from P22 Δgp16-infected cells, yielding 26,014 extended tail particles at 4.20 Å and a final set of 3,984 channel particles for reconstruction at 8.83 Å resolution.

### Model building and refinement

Previously determined structures of gp9 (PDB: 8EAN) and gp10 (PDB: 8EAP), together with AlphaFold3-predicted structures of gp7* and gp20, were used as initial templates for model building. These models were docked into the cryo-EM density maps and manually adjusted in Coot(*56*). The resulting models were refined using Namdinator(*57*) and phenix.real_space_refine(*58*) to optimize the fit to the experimental density. Statistics of the structural modeling are summarized in **Table S2**.

## Supplementary Materials

**Figure S1.**
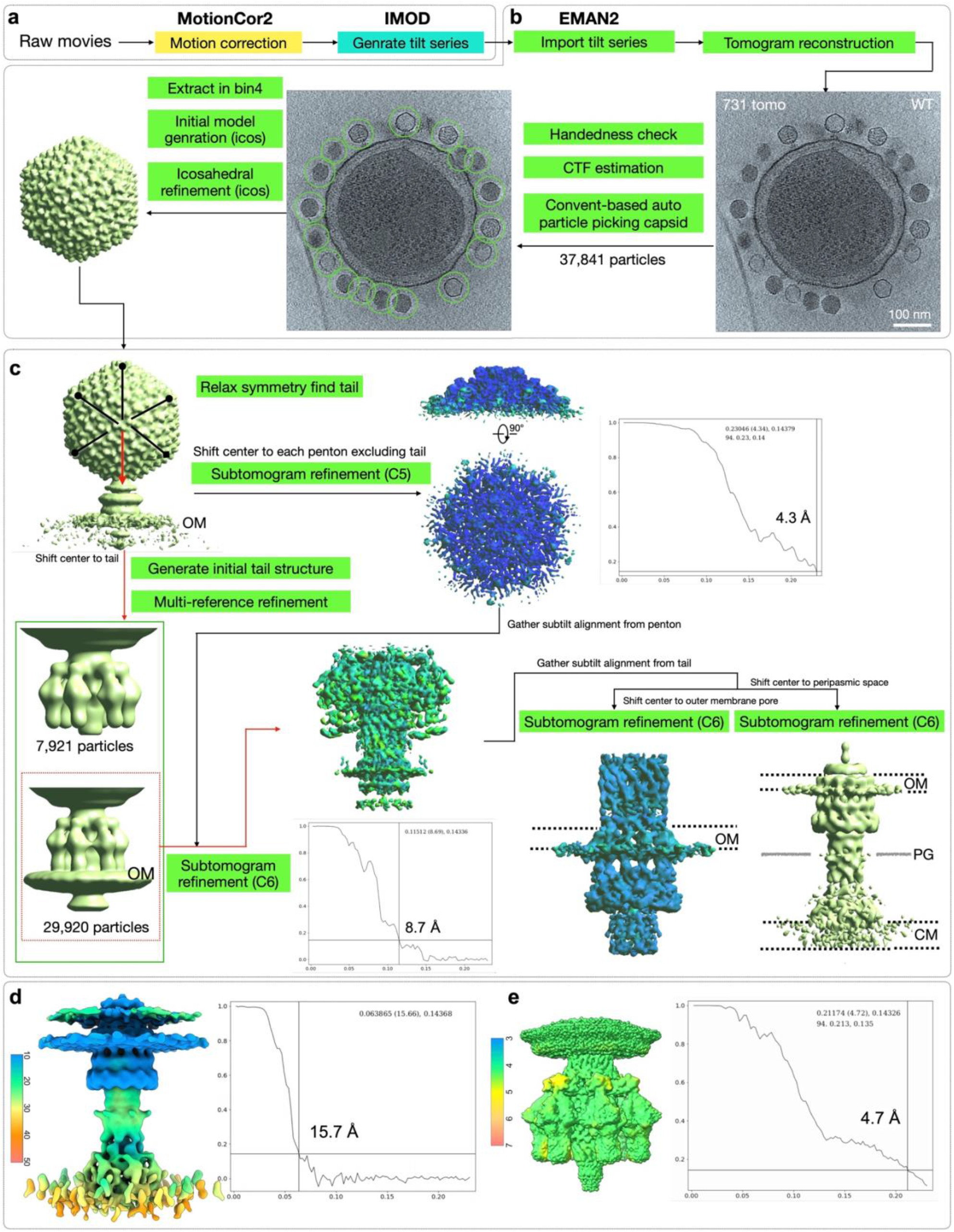
Cryo-ET data processing and structural determination of P22 during infection. (**a**) Preprocessing to generate tilt series. Raw movie frames were aligned using MotionCor2, followed by tilt-series stacking and alignment in IMOD. (**b**) Particle picking and averaging. Tomograms reconstructed in EMAN2 underwent per-tilt CTF estimation. A convolutional neural network-based auto-picking model, initially trained on ∼20 manually curated tomograms, achieved >90% accuracy across all 731 tomograms after iterative training and testing. Extracted capsid particles (binned by a factor of 4) were used for initial icosahedral, symmetry-guided, subtomogram averaging. (**c**) Local averaging of the extended tail and trans-envelope channel. Icosahedral symmetry was applied to refine well-defined capsids; symmetry was then released to locate tail vertices. C5-symmetrized vertex particles (4.3 Å resolution, Nyquist-limited) provided sub-tilt alignment parameters for downstream analysis that was applied to later steps. Tail vertices were extracted to generate an initial capsid-tail model, followed by multi-reference refinement to exclude incomplete assemblies. From 29,920 complete particles, iterative subtomogram refinement resolved the tail structure at 8.7 Å. Subsequently, tail sub-tilt alignment parameters were applied for subtomogram refinement of the membrane outer complex and periplasmic channel. (**d**) Local resolution estimation of the subtomogram-averaged periplasmic channel was performed using ResMap(*59*); the FSC curve of the periplasmic channel map from EMAN2 shows an overall resolution of 15.7 Å at the 0.143 cutoff. (**e**) Local resolution estimation and FSC curve of the extracellular complex of Δgp(16,20) averaged by EMAN2 shows an overall resolution of 4.7 Å at the 0.143 cutoff.

**Figure S2.**
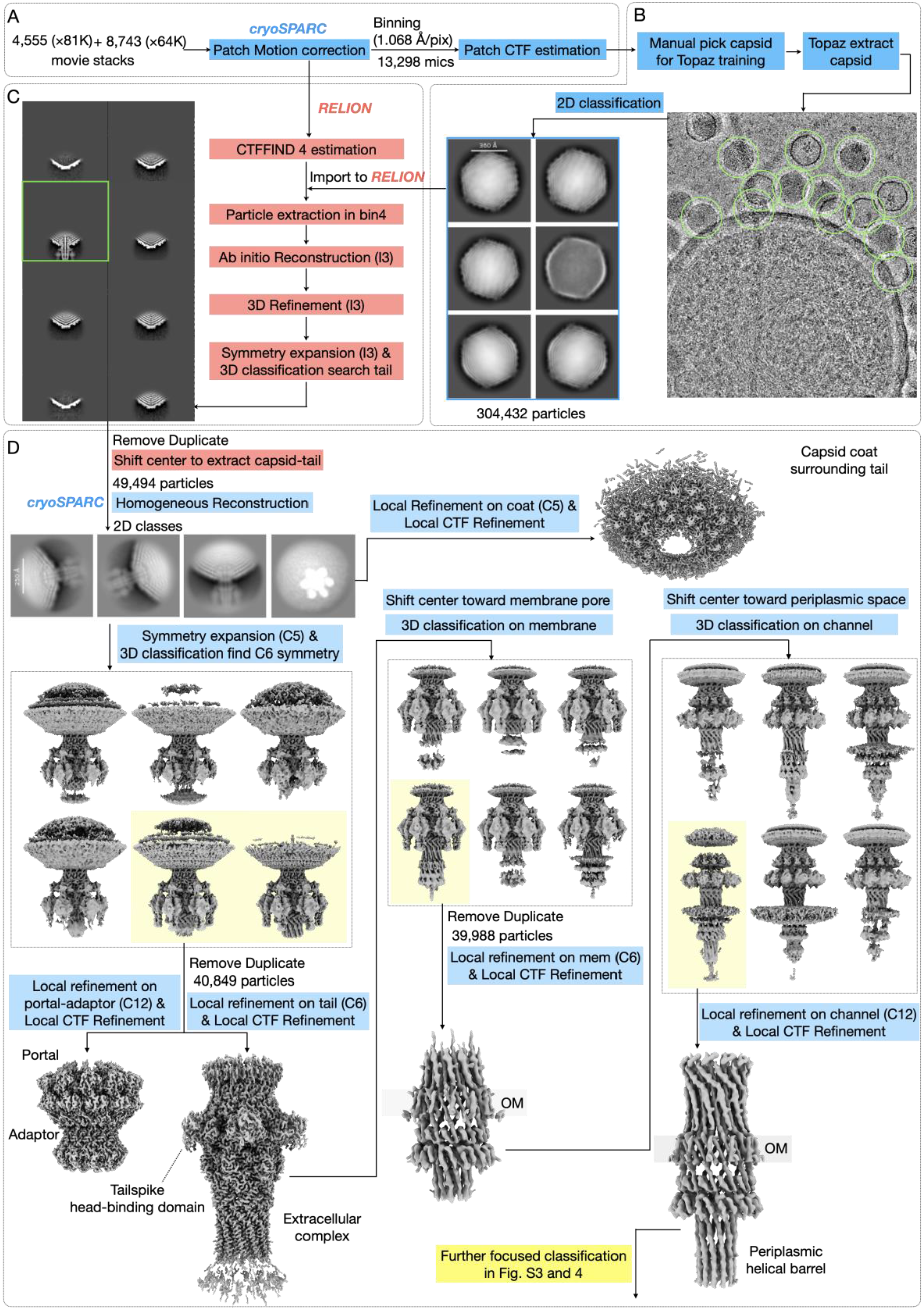
Cryo-EM data processing. (**a**) Pre-processing of cryo-EM images. A total of 8,743 movie stacks collected at ×64,000 magnification were motion-corrected in super resolution mode (0.673 Å/pixel) using Patch-Motion in cryoSPARC and subsequently cropped to a final pixel size of 1.068 Å using the *relion_image_handler*, while 4,555 movie stacks collected at ×81,000 magnification underwent Patch-Motion correction in cryoSPARC with a binning factor of 2 (final pixel size: 1.068Å/pixel). All micrographs, uniformly pre-processed to the same pixel size, were subjected to Patch-CTF estimation in cryoSPARC. (**b**) Particle picking and initial screening. Capsids were picked using a trained Topaz model based on a manually curated template, followed by 2D classification in a preliminary screen for particle quality. This yielded 304,432 particles, which were extracted and imported into RELION for further analysis. (**c**) Global localization of the tail. Particles imported from cryoSPARC into RELION were re-extracted in RELION at a binning factor of 4, and 3D refinement oriented to I3 icosahedral symmetry. After capsid refinement the tail was localized by applying 60-fold symmetry expansion, followed by iterative 3D classification without image alignment. This approach generated symmetry-related orientations for each particle, enabling global angular searching to align and classify tail structure. Classes exhibiting well-defined tail vertices were prioritized for downstream processing. (**d**) Focused refinement of the tail and trans-envelope channel. Duplicate particles were removed and their box centers were shifted to align with the corresponding tail complex. The capsid shell underwent direct focused refinement using C5 symmetry, while symmetry expansion combined with masked 3D classification was employed to resolve the phage tail structure using C6–C12 symmetry. This step included local refinement of the extracellular complex, portal-adaptor complex, outer membrane complex, and periplasmic channel.

**Figure S3.**
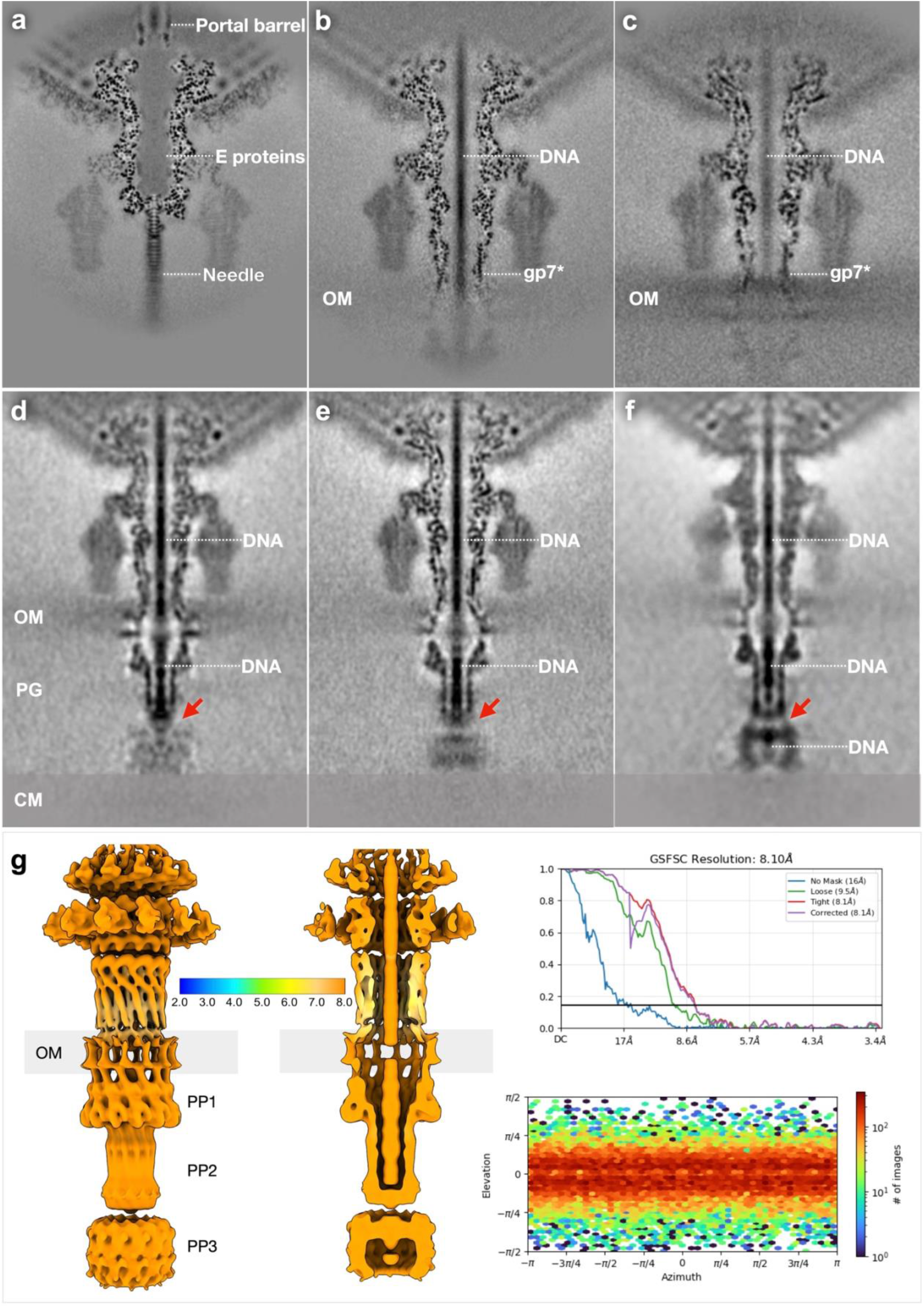
Comparison of mature and infecting P22 particles. (**a**) Tail complex of mature wild-type P22; E proteins are primarily stored within the portal and tail lumina. Adapted from EMD-27790. (**b, c**) Extracellular complexes of wild-type (**b**) and Δgp16 (**c**) during infection. (**d–f**) Classification of DNA-filled phage particles revealed three structural states of the periplasmic channel. These states potentially correspond to sequential infection intermediates, with the C-terminal helical barrel of gp20, indicated by arrows, appearing to progressively open where it connects to its “club-foot”. N indicates the particle number in each class. Only in panel F does DNA density extend all the way to the cytoplasmic membrane. (**g**) Structural evaluation of the trans-envelope channel.

**Figure S4.**
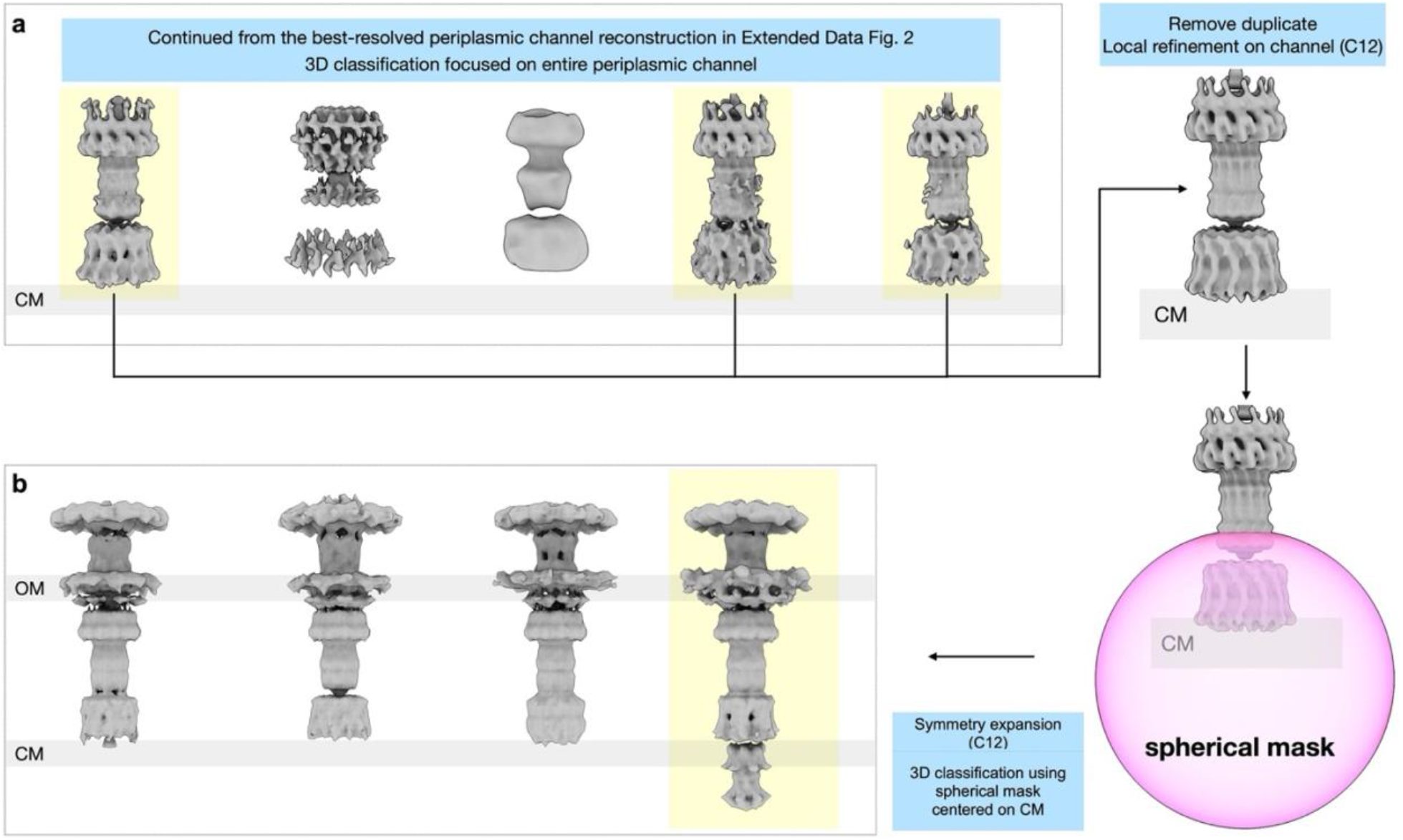
Cryo-EM data processing of the P22^+^ gp20 PP3 domain and gp16. (**a**) Following symmetry expansion and 3D classification, particles with a well-resolved gp20 PP3 domain were subjected to local refinement. Focused classification was then performed using a spherical mask centered on the cytoplasmic membrane (CM). (**b**) Four representative classes were resolved, one of which contains an additional density spanning the CM. This class was not seen in a comparable focused classification of Δgp16-infected minicells, allowing us to conclude that the density spanning the CM is gp16.

**Figure S5.**
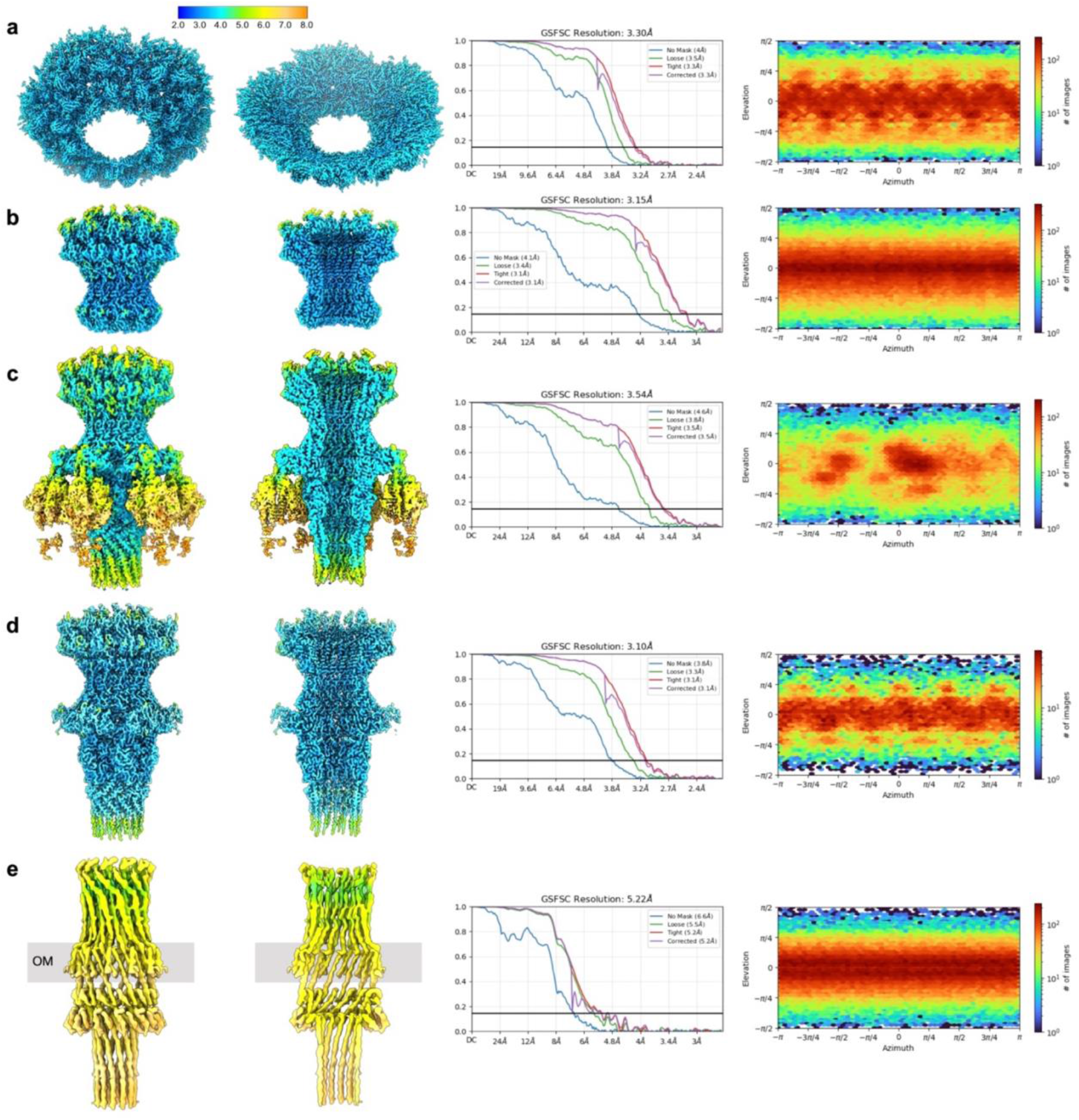
Cryo-EM structure evaluations. Local resolution estimation map, Fourier shell correction (FSC) at 0.143 and Euler angle distribution of refined particle subset used in the final cryo-EM reconstruction of each substructure. (**a**) Capsid vertex surrounding the tail complex. (**b**) Portal-adaptor complex. (**c**) Extracellular complex including tailspikes. (**d**) Extracellular tail complex. (**e**) Extracellular channel, outer membrane and periplasmic channel without the PP3 domain of gp20.

**Figure S6.**
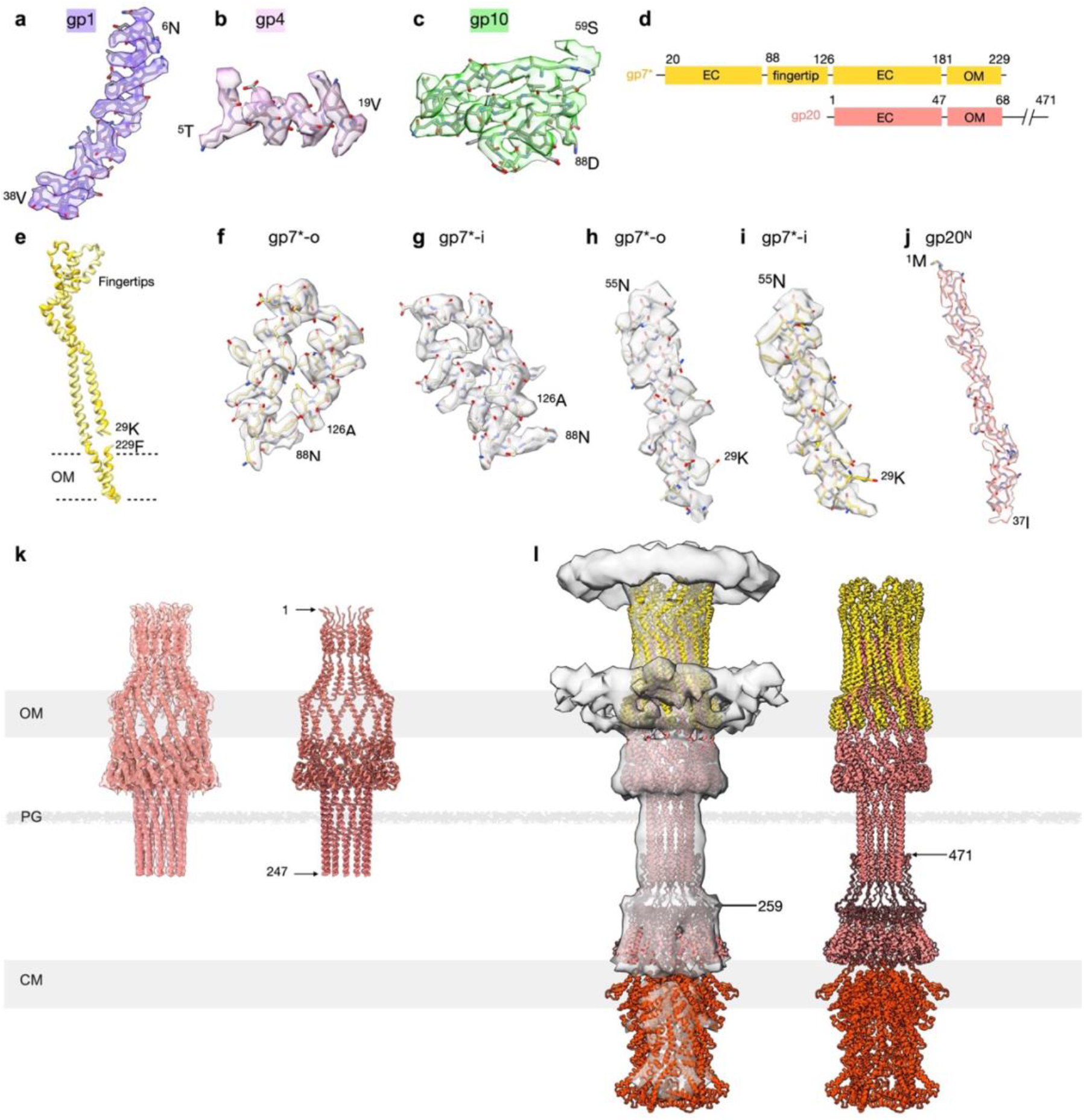
Superposition of cryo-EM densities and protein models. (**a–c**) Representative model fitting of gp1, gp4, and gp10. (**d**) Genome maps of extracellular domains of gp7* and gp20. (**e**) Superposition of gp7*-o (light yellow) and gp7*-i (yellow) shows that most regions align well, with differences restricted to the fingertip domains. (**f, g**) Model fitting of the fingertip domains of gp7* in the two conformations interacting with gp10. (**h, i**) Model fitting of the truncated N terminus of gp7*-o and gp7*-i adopting similar conformations. (**j**) Model fitting of extracellular domain of gp20. (**k**) *Left:* A truncated gp20 terminating at the helical barrel end (residue 247) fitted into the 5.22 Å cryo-EM density; *Right:* the fitted model alone. (**l**) *Left*: The entire trans-envelope channel structure, including the cryo-EM resolved models of gp7* and gp20 and the AlphaFold3-predicted PP3 domain of gp20 and gp16 fitted into the cryo-EM density; *Right*: the fitted model alone.

**Figure S7.**
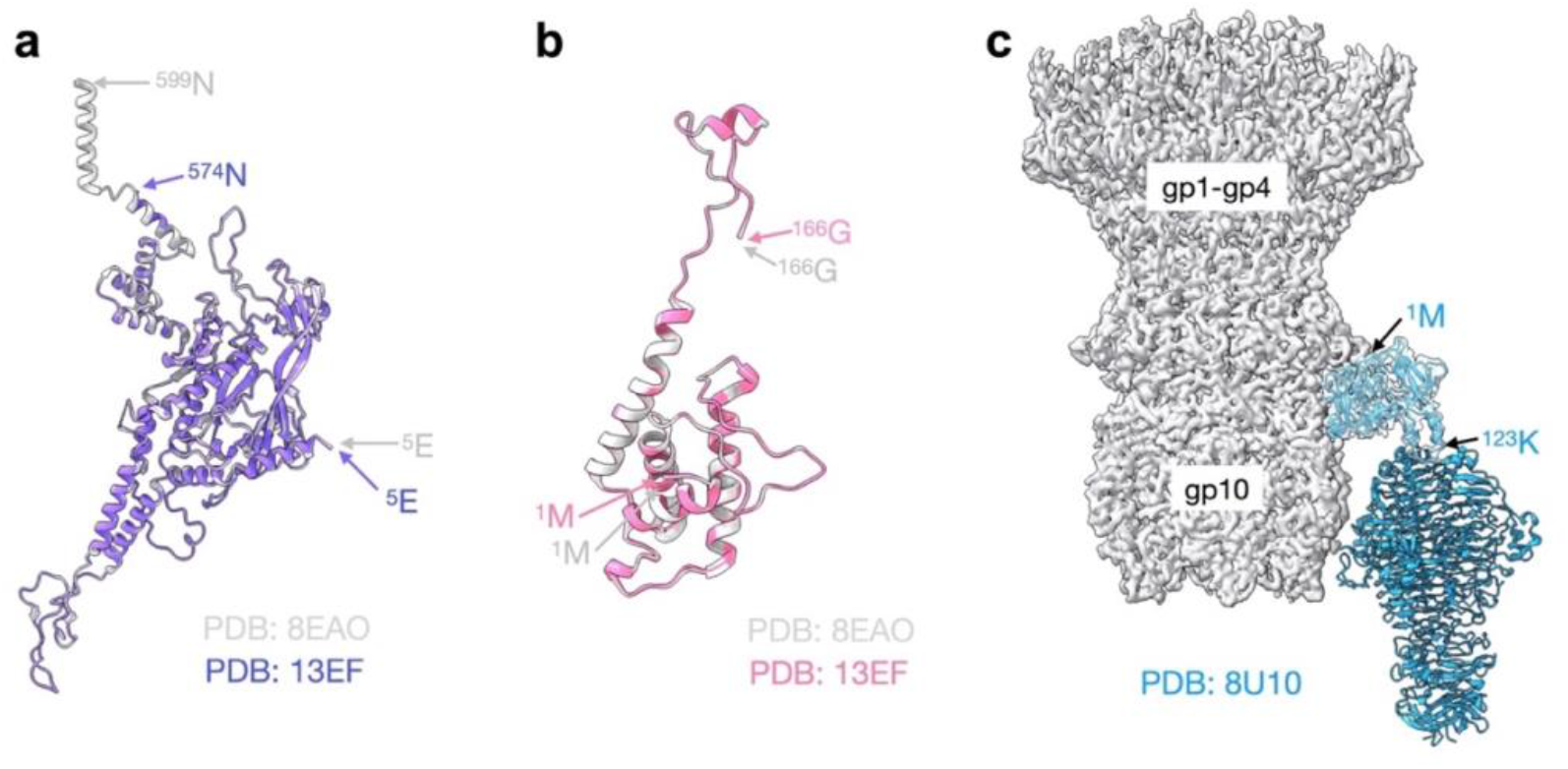
Structural comparisons before and after infection. (**a**) Portal, gp1 (Mature: PDB: 8EAO, gray; Infection: PDB: 13EF, purple). (**b**) Tail adaptor, gp4 (Mature: PDB: 8EAO, blue; Infection: PDB: 13EF, pink). (**c**) Tailspike protein gp9 from mature P22 (PDB: 8U10) fitted into the P22 density after infection. Only the head-binding region (^1^M**–**^123^K) of gp9 was clearly resolved at high resolution.

**Figure S8.**
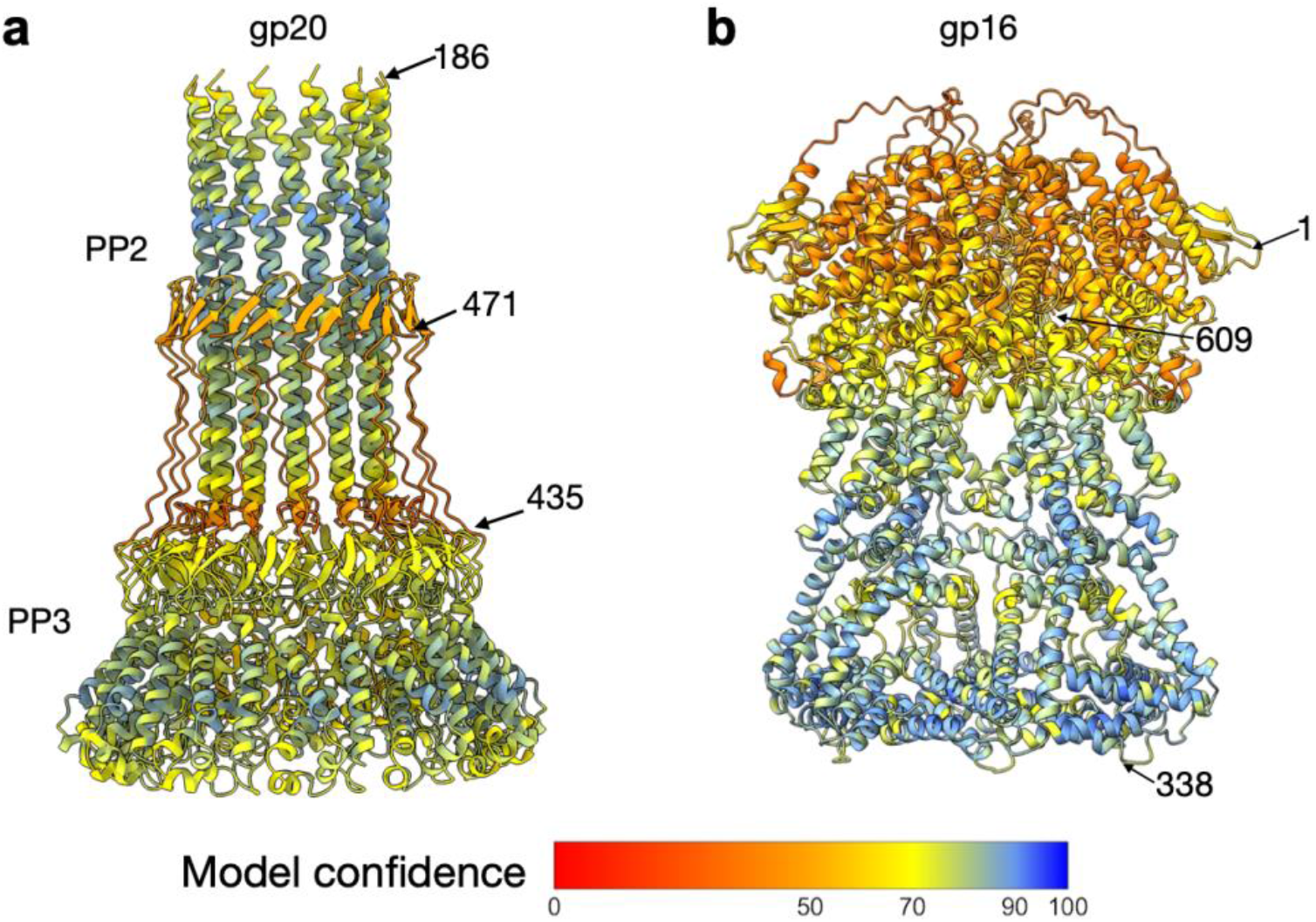
AlphaFold3-predicted structures of gp20 and gp16. (**a**) Side view of the predicted hexamer structure of the gp20 PP2–PP3 domains, residues 186–471. (**b**) Side view of the predicted structure of the gp16 hexamer.

**Figure S9.**
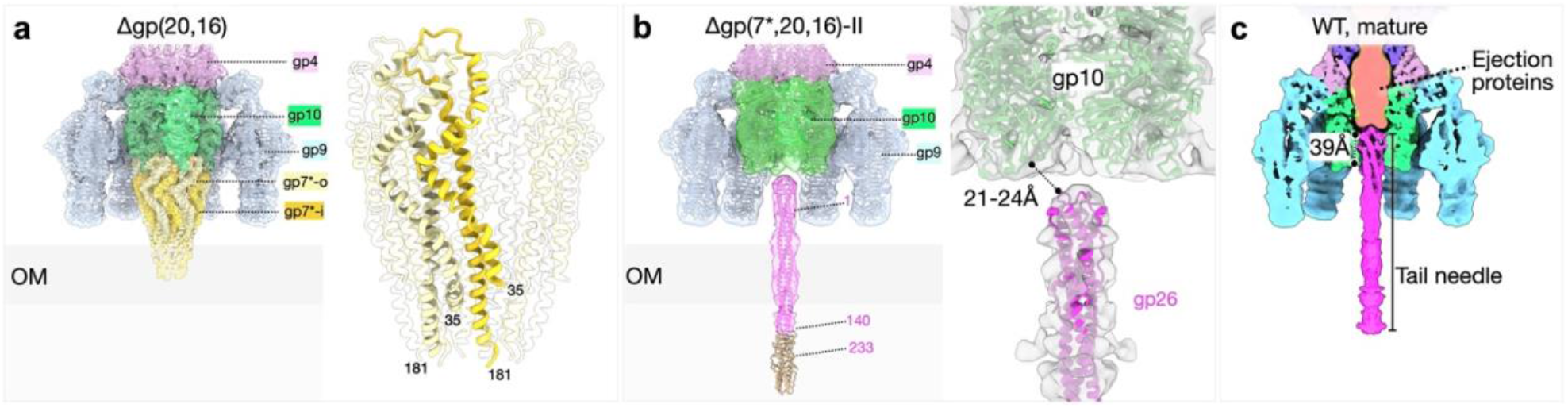
Structural comparison of mutant and wild-type phage. **(a)** *Left:* Subtomogram-averaged structure of the Δgp(16,20) tail with the pseudo-atomic model of P22^+^ fitted into the density. *Right:* A close-up view shows the gp7* dodecamer, residues 35–181 of gp7*-o and gp7*-i are resolved. **(b)** *Left:* Subtomogram-averaged structure of the Δgp(7*,16,20)-II tail with the P22^+^ pseudo-atomic model fitted into the cryo-ET density. The density corresponding to the disengaged gp26 tail needle (residues 1–140, in magenta) was fitted using PDB: 2POH, whereas the C-terminal portion of the needle was not resolved. The remaining tail components were modeled using gp4– gp10 (PDB: 13EF) and gp9 (PDB: 8EAN). *Right:* Zoom-in view of the tail hub and fitted gp26 needle, showing that the gap between gp26 and the gp10 hub exceeds ∼20 Å. (**c**) Structure of a mature P22^+^ virion showing the gp26 tail needle inserted approximately 39 Å into the gp10 tail hub (adapted from EMD-27790 and 27792).

**Figure S10.**
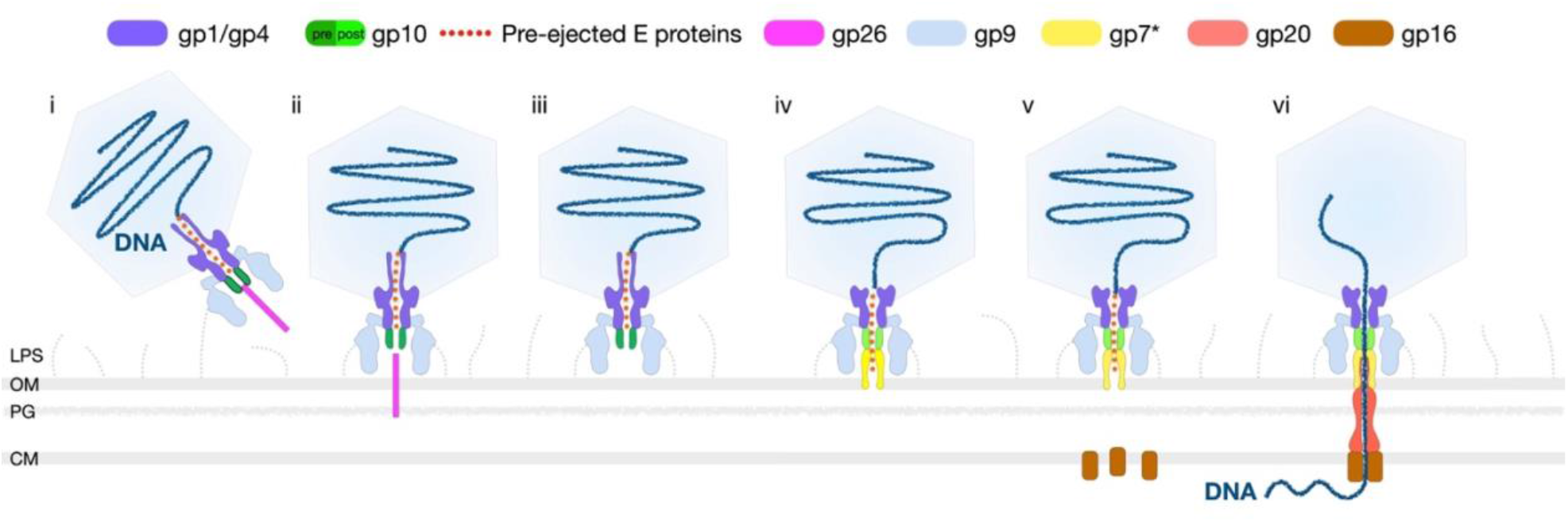
Schematic models of wild-type P22 infection. (i) P22 adsorbs to a cell, where tailspikes degrade O antigen and bring the particle close to the OUTER MEMBRANE surface. Because the tail needle extends below the plane of the tailspikes, P22 initially interacts with the cell surface at an angle but (ii) continued hydrolysis of O antigen reorients the particle perpendicular to the outer membrane, with the tail needle penetrating the OUTER MEMBRANE. (iii) A small conformational change in the gp10 hub triggers needle release, which in turn leads to (iv) ejection of 12 copies of gp7*; these assemble into an extracellular channel that enters the outer membrane. (v) Gp16 is ejected and passes through the gp7* channel into the cell and localizes to the CYTOPLASMIC MEMBRANE. (vi) Finally, gp20 is ejected; it stabilizes the gp7* structure in the outer membrane and also forms a channel-like structure through the PG and across the periplasm where it can interact with gp16 to complete the trans-envelope channel necessary for DNA ejection into the cytoplasm.

**Figure S11.**
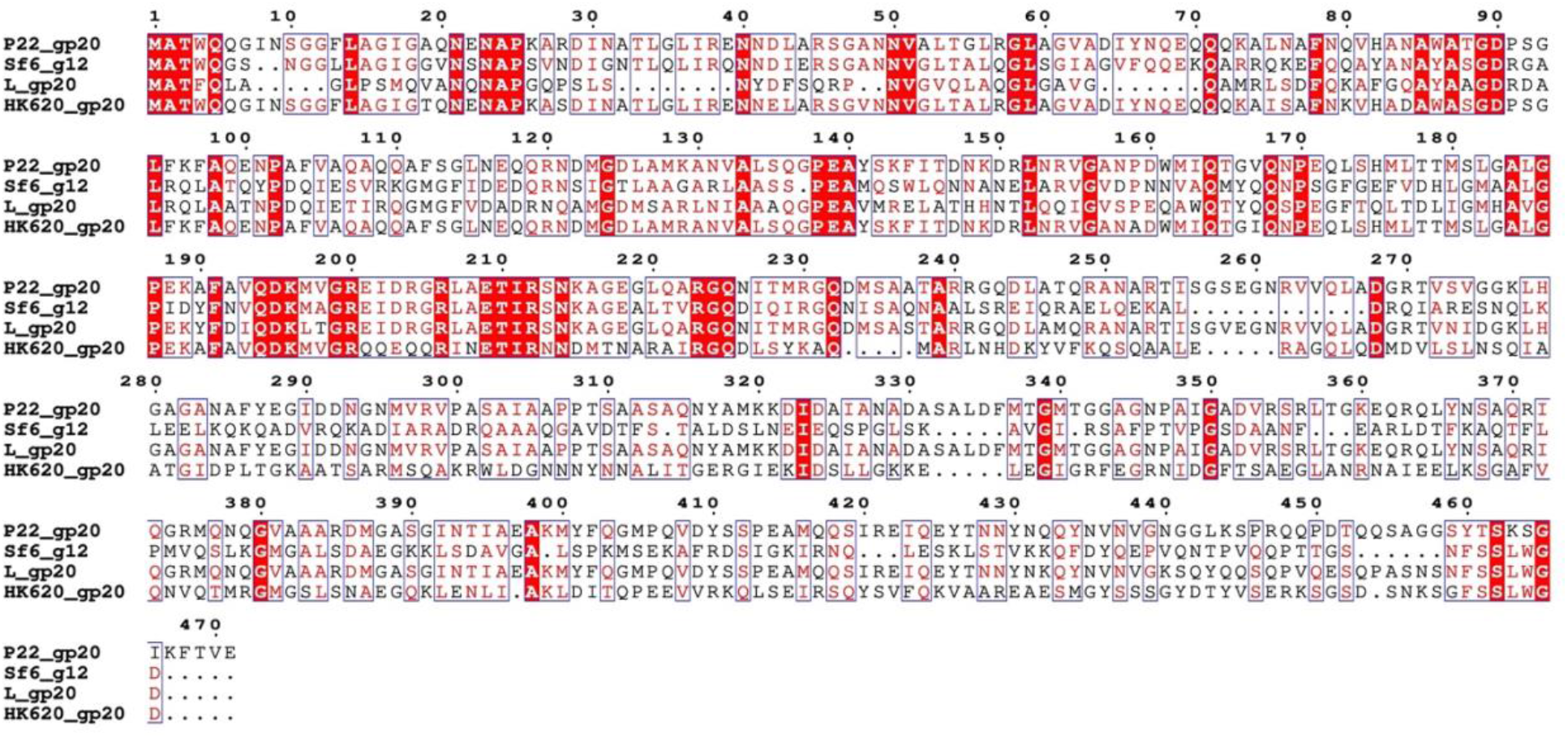
Sequence alignment of P22 gp20. Identities to P22 gp20 (residues 1–259) are 46.4% (Sf6), 47.3% (L) and 78.5% (HK620).

**Table S1.**
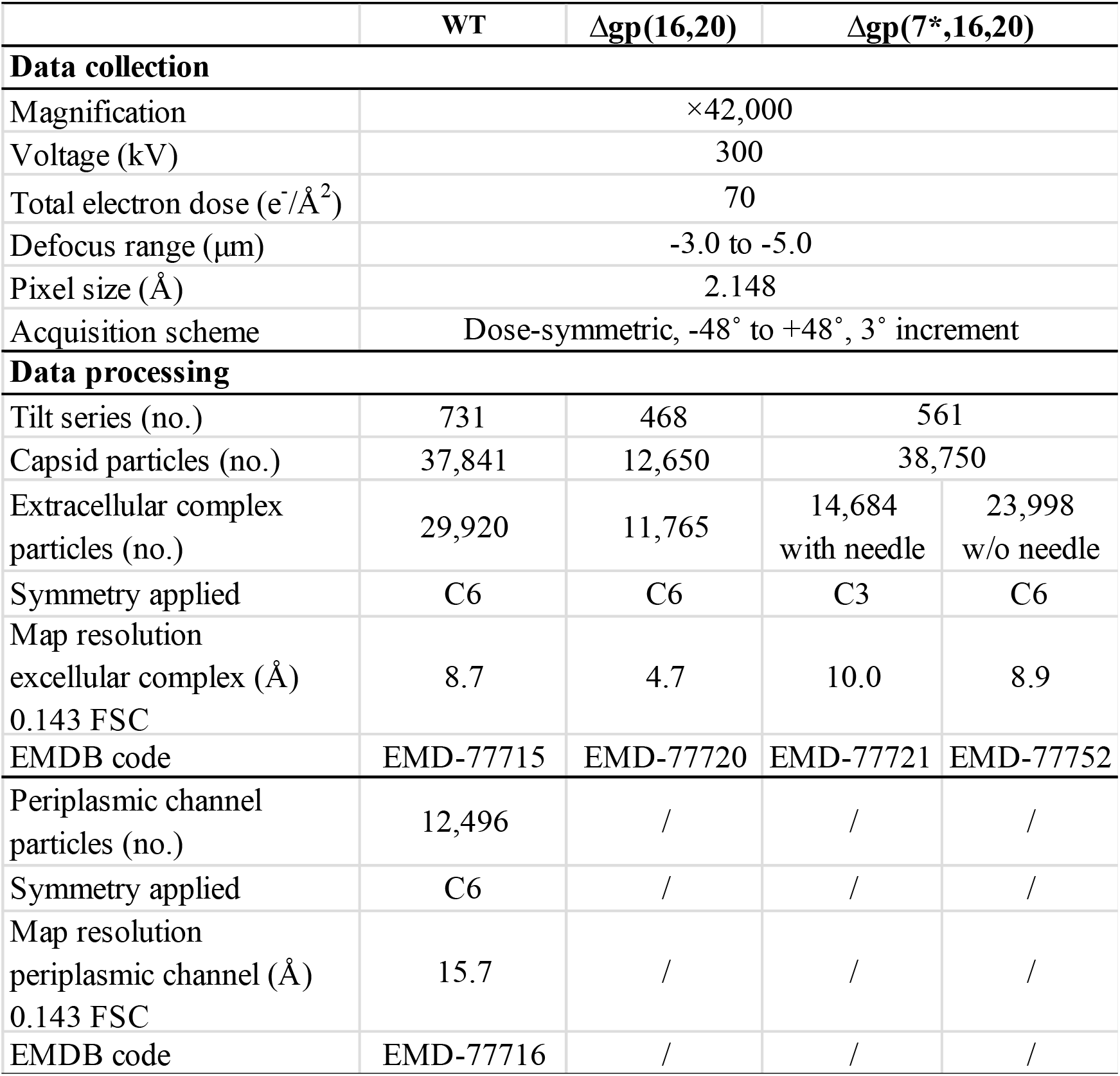
Cryo-ET data collection and processing statistics.

**Table S2.** Cryo-EM data collection, processing and structure refinement statistics.

| | Excellular<br>channel (WT) | OM and periplasmic<br>channel (WT) | Periplasmic and<br>cytoplasmic<br>channel (WT) | Overall<br>trans-envelope<br>channel (Δgp16) |
| --- | --- | --- | --- | --- |
| Data collection |  |  |  |  |
| Magnification | $\times 81,000/\times 64,000$ | | $\times 64,000$ | $\times 81,000$ |
| Voltage (kV) | 300 |  | 300 | 300 |
| Total electron dose ($e^-/\text{\AA}^2$) | 55 | | 55 | 55 |
| Defocus range ($\mu\text{m}$) | -1.6 to -2.2 | | -1.6 to -2.2 | -1.6 to -2.2 |
| Pixel size ($\text{\AA}$) | 1.068 | | 1.346 | 1.068 |
| Micrographs (no.) | 13,298 |  | 4,153 | 10,453 |
| Data processing |  |  |  |  |
| Symmetry imposed | C6 | C12 | C6 | C6 |
| Final particle images (no. ) | 119,473 | 27,985 | 5,406 | 29,180 |
| Map resolution ($\text{\AA}$) 0.143 FSC | 3.10 | 5.22 | 10.61 | 7.29 |
| Map sharpening B factor ($\text{\AA}^2$) | -77.6 | -236.7 | / | -360.0 |
| Refinement |  |  |  |  |
| CC (model vs. data) | 0.82 | 0.8 | / | / |
| Chain count | 72 | 24 | / | / |
| Non-hydrogen atoms | 122,898 | 40,776 | / | / |
| Protein residues | 15,642 | 5,550 | / | / |
| Ligands | 0 | 0 | / | / |
| B factors ($\text{\AA}^2$) | | | | |
| Proteins | 88.23 | 135.54 | / | / |
| Ligands | / | / | / | / |
| R.m.s. deviations |  |  |  |  |
| Bond lengths ($\text{\AA}$) | 0.003(0) | 0.008 (0) | / | / |
| Bond angles ($^\circ$) | 0.603(172) | 1.068 (37) | / | / |
| Validation |  |  |  |  |
| MolProbity score | 2.08 | 2.77 | / | / |
| Clashscore | 8.96 | 7.51 | / | / |
| Ramachandran plot |  |  |  |  |
| Favoured (%) | 95.43 | 92.17 | / | / |
| Allowed (%) | 4.52 | 6.49 | / | / |
| Outliers (%) | 0.05 | 1.34 | / | / |
| PDB code | 13EF | 12BO | / | / |
| EMDB code | EMD-77019 | EMD-76290 | EMD-78681 | EMD-77764 |

**Movie S1.** Animation showing *in situ* structures derived from cryo-ET/subtomogram averaging and *in situ* single-particle cryo-EM.

**Movie S2.** Animation showing wild-type P22 infection initiation.

**Movie S3.** Animation showing extra-cytoplasmic complementation by coinfecting Δgp20 and Δgp16. For simplicity, only six copies of gp16 are modeled

